# Bacteriophages utilize pseudolysogeny to target non-replicating bacteria and CRISPR-resistant phages eliminate recalcitrant implant infections

**DOI:** 10.64898/2026.03.24.714066

**Authors:** Yeswanth Chakravarthy Kalapala, Aditya Kamath Ammembal, Saksham Jain, Nisha Sanjay Barge, Rachit Agarwal

## Abstract

A key driver of bacterial infection treatment failure and relapse is the persistence of non-replicating bacterial subpopulations that emerge under stressors like nutrient starvation and immune pressure. These dormant cells evade antibiotics, fuelling recurrence and resistance. Bacteriophage therapy is a promising alternative, but its efficacy against non-replicating bacteria is poorly understood. Improving our understanding of bacteria-phage interactions under non-replicating conditions could greatly enhance phage therapeutic outcomes in clinics. By utilising various bacterial (*Mycobacterium smegmatis*, *Mycobacterium tuberculosis*, and *Pseudomonas aeruginosa*) and phage species, this study quantitatively demonstrates that lytic phages can infect non-replicating bacteria (under nutrient starvation, acidic pH or antibiotic pressure), persisting in a state of pseudolysogeny and resuming lysis upon bacterial regrowth. We find that the pseudolysogeny window is phage- and host-dependent, with degradation of extrachromosomal phage DNA leading to loss of pseudolysogeny. We find that Pseudomonas CRISPR defence plays a crucial role in phage DNA degradation even under non-replicating conditions, underscoring the need for its consideration in phage therapy design. We also demonstrated the *in vivo* relevance of pseudolysogeny and CRISPR-resistant bacteriophages in eliminating implant-associated and antibiotic-persistent Pseudomonas infections in mice. These findings highlight the need to consider phage-host dynamics and bacterial defences when designing phage-based strategies to target non-replicating bacteria and persistent infections.

## INTRODUCTION

Bacterial infections cause 1 in 8 deaths globally (13.6% of total deaths), accounting for 7.7 million fatalities annually^1^. Antibiotic treatments are under pressure from the emergence of antimicrobial resistance (AMR) and a dwindling antibiotic pipeline^2^. The efficacy of antibiotics hinges on bacterial physiology and is compromised by population heterogeneity^3^. Non-replicating and slow-metabolising tolerant bacteria are selected in patients exposed to antibiotics^4^. Once antibiotics are cleared from the body, these slow-growing or dormant cells have been shown to drive relapse in infections associated with *Staphylococcus aureus*^5^, *Pseudomonas aeruginosa*^6^, and *Mycobacterium tuberculosis*^7^, despite completion of standard treatments.

Triggered persistence or tolerance in bacteria can arise from nutrient deprivation in the bacterial microenvironment. Intracellular pathogens such as *M. tuberculosis*^8^ and *Salmonella spp.*^9^ enter a state of persistence due to host-associated factors and the host immune system. Ironically, antibiotics themselves trigger persistence in several bacterial species^3^. These antibiotic-induced persisters often survive subsequent exposure to very high concentrations of antibiotics even in the absence of heritable mutations or antibiotic resistance genes^3^.

Amid this global crisis, lytic bacteriophage-based therapies are emerging as a promising alternative^10^. Bacteriophages are viruses that are highly selective, potent, and exhibit minimal immunogenicity^10,11^. Their efficacy against both antibiotic-sensitive and antibiotic-resistant bacteria is well documented^10^. Lytic phage therapy has been compassionately administered for numerous antibiotic-resistant bacterial infections worldwide^12–14^. Treatment of recurrent infections with phage cocktails along with antibiotics has been shown to significantly reduce relapse rates across several bacterial species, including *K. pneumoniae*^15^, *P. aeruginosa*^16,17^ and *E. coli*^18^. For instance, a study investigating phage-antibiotic combination therapy for periprosthetic hip joint infections demonstrated an eightfold reduction in relapse rates compared to antibiotics alone^19^. While these findings underscore the potential of lytic phage therapy to eradicate infections and lower relapse rates, emerging evidence highlights the growing complexity of bacteriophage lytic lifecycles, with phage-carrier states and pseudolysogeny representing important alternative infection strategies^20^. Despite promising outcomes in compassionate use cases, poorly understood phage-host interactions and the emergence of phage resistance remain significant barriers to the adoption of phage therapy as a mainstream treatment for bacterial infections. Identifying the critical gaps in our mechanistic understanding of phage efficacy against bacterial subpopulations, especially persistent bacterial subpopulations could help us exploit the full spectrum of phage lifecycles and further reduce the relapse rates.

Studies conducted in stationary-phase bacterial cultures have shown promising results. Bacteriophage T4 effectively infects and lyses stationary-phase *Escherichia coli*^21^. Similarly, SEP1 virus showed infection dynamics capable of targeting stationary-phase *Staphylococcus epidermidis*^22^. In our previous work, mycobacteriophages D29, TM4, and Che7 demonstrated the ability to lyse stationary-phase *Mycobacterium smegmatis*^23^. However, stationary-phase models represent a heterogeneous bacterial state encompassing replicating, dying, and dormant cells, thereby inadequately recapitulating true bacterial dormancy^24^. Several studies have demonstrated the ability of bacteria to grow in the stationary phase through the recycling of nutrients from dead bacteria^25,26^. Therefore, stringent stress models where bacteria enter a state of non-replication in response to nutrient-starvation, acidic pH and antibiotics, provide a more translatable framework to study bacterial tolerance and persistence. For example, investigations into *Lactobacillus plantarum* bacteriophage B1 revealed that phage replication and cell lysis were suppressed under nutrient-deprived conditions, with phage release occurring only upon restoration of favourable growth conditions^27^. Similar observations in other lytic phage-host systems suggest a phenomenon of pseudolysogeny, wherein bacteriophages remain inactive as extra-chromosomal DNA within nutrient-deprived host cells until host metabolism resumes, enabling lytic transition^28–30^. Recent widespread detection of lytic phage genomes across more than 1,000 species and millions of genomes highlights phage pseudolysogeny as a pervasive component of phage biology^31,32^.

Yet fundamental questions remain unanswered. While some phages can infect bacteria in the stationary phase, the infectivity of non-replicating bacteria, characterised by significant cell wall remodelling and receptor downregulation^33,34^, remains largely unexplored. Additionally, the stability of non-integrating lytic phage DNA within non-replicating cells during pseudolysogeny and its potential for reactivation upon host resuscitation remain unclear.

Furthermore, the status of anti-phage defence, including non-specific defences and CRISPR-Cas systems in non-replicating cells warrants investigation. Although previous studies have shown the activity of CRISPR-Cas systems under bacteriostatic conditions^35^, it remains unclear how this affects the pseudolysogeny window of bacteriophages. Unravelling this black box of bacteriophage-bacteria interactions under non-replicating conditions could hold the key to more rational bacteriophage engineering strategies for treating persistent bacterial populations and preventing relapses.

In this study, we addressed these gaps by systematically analysing bacteriophage-host interactions under non-replicating conditions induced by nutrient deprivation, antibiotic pressure or low pH by using lytic bacteriophages. Specifically, we investigated the infectivity, persistence, and reactivation of bacteriophages within non-replicating bacteria and the functional status of bacterial immune systems in these states. Key results were validated using single cell imaging and analysis. We also demonstrated the *in vivo* relevance of pseudolysogeny in a mouse model of implant-associated Pseudomonas infection. Notably, we demonstrated for the first time that CRISPR-resistant bacteriophages completely eliminated all bacteria in an antibiotic-persistent, implant-infection model in mice and outperformed CRISPR-susceptible bacteriophages. Understanding the dynamics of phage infection in non-replicating systems is critical to advancing phage-based strategies for combating persistent infections, offering a novel avenue to tackle the escalating challenge of AMR.

## RESULTS

### 1. Phages infected non-replicating *M. smegmatis*

Bacteria are observed to undergo significant cell wall remodelling when entering a state of non-replicating persistence^33,34^. This cell wall remodelling could downregulate some of the key receptors required by bacteriophages for successful infection. We began by establishing a non-replicating *M. smegmatis* model where log-phase cultures of *M. smegmatis* were starved by resuspending the culture in Phosphate Buffered Saline (PBS) for 48 h. We confirmed that a 48 h starvation period is successful in driving the surviving bacteria into a state of non-replication by enumerating the clock plasmid loss^36^ over prolonged periods of time **(Fig 1A, Supp Fig 1A, B)**. We also checked for the upregulation of key non-replication markers in bacteria^37,38^ – relA and acyl-coA synthase, which showed an upregulated expression **(Supp Fig 1C)**. Using this non-replicating model for *M. smegmatis,* we checked for phage infectivity by exposing the 48 h starved bacteria to GFP encoding □^2^GFP10 bacteriophage (derived from TM4) at 10:1 Multiplicity of Infection (MOI) for different periods of time **(Fig 1B)**. We observed that over 50% of the bacteria were infected with a 30 min exposure to □^2^GFP10, and the infection maxima was at around 3 h, where 80% of the total bacterial cells were infected with the bacteriophage. We further validated phage infectivity of non-replicating *M. smegmatis* by SYTO9-stained TM4 bacteriophage and found 70% of *M. smegmatis* showing SYTO9 signal 3 h post-infection **(Supp Fig 2)**. Infection efficacy of □^2^GFP10 against actively replicating bacteria has been reported to be around 72% at MOI of 10:1^39^. These findings are also in line with previous evidence of phages infecting non-replicating bacteria in other bacterial systems^21,22,40,41^, and for the first time, provide quantitative insights into mycobacteriophage efficiencies in infecting non-replicating bacteria.

**Figure 1.**
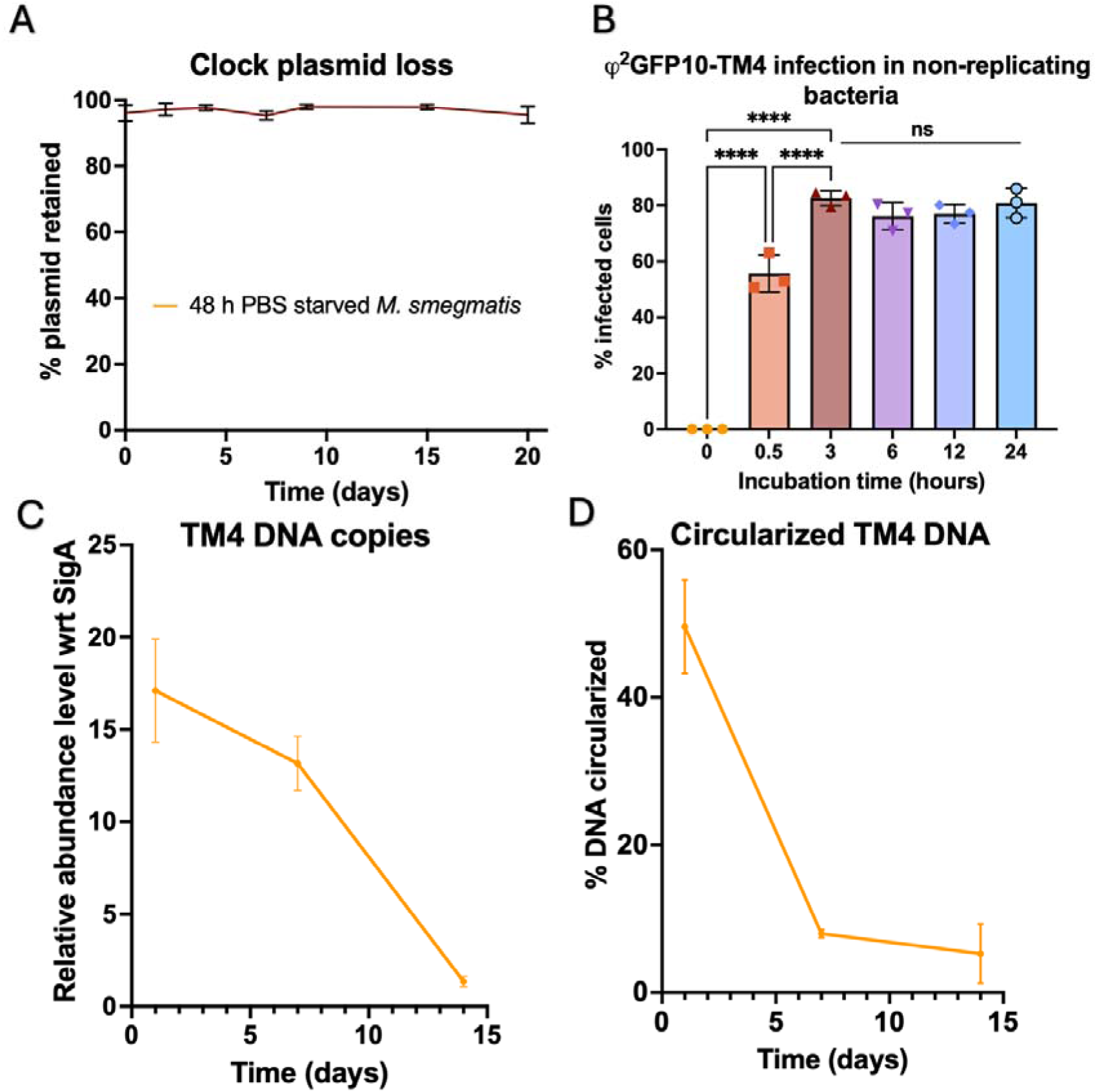
Phage infection in non-replicating M. smegmatis. **A)** Percentage of surviving bacteria with the clock plasmid retained post 48 h PBS starvation (n=3) **B)** Phage infection towards non-replicating M. smegmatis validation using □^2^GFP10 bacteriophage at 37°C (n=3). One-way ANOVA with multiple comparisons was used to estimate the statistical significance. (****p ≤0.0001). **C)** TM4 Phage DNA quantification (n=3). **D)** Circularization estimation in non-replicating M. smegmatis post phage exposure (n=3).

**Figure 2.**
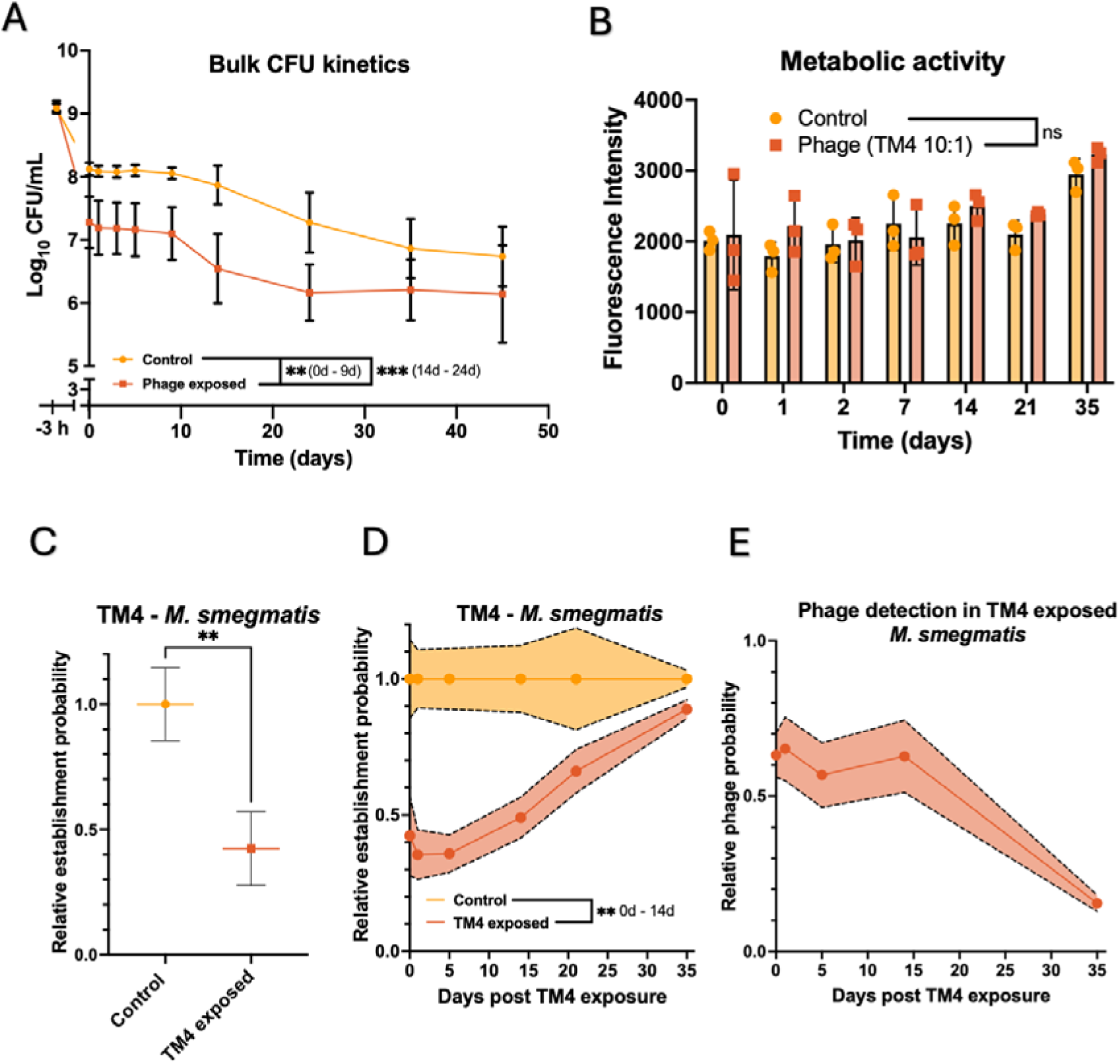
Phage pseudolysogeny mediated killing of non-replicating bacteria. **A)** Bulk kinetics of 48 h PBS starved M. smegmatis exposed to TM4 bacteriophage at 10:1 MOI. Negative timepoint is obtained prior to treatments and immediately post starvation (n= 5). **B)** Resazurin assay of 48 h PBS starved M. smegmatis cells in PBS from Day 0 up to Day 35 (n=3). **C)** TM4 phage infectivity and efficacy towards non-replicating M. smegmatis quantified through single-cell growth assay (n=3). **D)** TM4 pseudolysogeny window in non-replicating M. smegmatis quantified through single-cell growth assay (n=3). **E)** Probability of phages in non-replicating M. smegmatis quantified through single-cell growth assay (n=3). For A, multiple unpaired t-tests were done and for B and D, two-way ANOVA with Holm-Šídák’s multiple comparisons test was used to estimate the statistical significance between groups. For C, unpaired t-test was used to determine the statistical significance (*p ≤ 0.05).

### 2. Phage DNA remained detectable and viable for several days

To further validate the phage infection of non-replicating *M. smegmatis*, we extracted the total DNA from phage-exposed non-replicating *M. smegmatis* samples post viricide (Ammonium Ferric Sulphate) treatment and 3X PBS washes and performed quantitative PCR to detect phage DNA with phage-specific primer pair. We were able to detect around 10-20 DNA copies of TM4 bacteriophage DNA per bacterial copy and the TM4 DNA copy numbers remained stable over a period of 7 days **(Fig 1C)** beyond which the phage DNA starts diminishing. Phage circularization is a crucial process for replication in several phages, including DNA mycobacteriophages^42^, such as TM4, and determines the stability of phage DNA against non-specific exonuclease degradation^43,44^. Quantitative PCR to detect the circularization of TM4 revealed around 50% circularization of phage DNA at day 0 in non-replicating *M. smegmatis* **(Fig 1D)**. Remarkably, we noticed the circularization detection to also decrease with prolonged pseudolysogeny, indicating phage DNA degradation and loss of pseudolysogeny.

### 3. Phages lysed non-replicating *M. smegmatis* upon revival

Having confirmed phage infectivity against non-replicating *M. smegmatis* and phage DNA stability, we proceeded to assess the phage efficacy in lysing PBS-induced non-replicating *M. smegmatis*. We exposed non-replicating *M. smegmatis* to TM4 at an MOI of 10:1 for 3 h and washed the cells three times with PBS and once with Ammonium Ferric Sulphate (AFS) for 5 mins to remove and/or inactivate extracellular phages^45^. The cultures were incubated in PBS at 37 °C at 180 RPM, and at periodic intervals, cultures were aliquoted, and the Colony Forming Units (CFUs) were enumerated on 7H10 growth medium. The starting CFU before phage treatment has been marked as −3 h time point. We observed around 80% reduction in bacterial CFU counts exposed to phages compared to non-phage exposed bacteria for the first 25 days **(Fig 2A)**. This would indicate that phage treatment resulted in either killing of the non-replicating bacteria in PBS or phage mediated killing of bacteria as it regrows on nutrient containing agar plates. Phage PFU in the PBS supernatant continued to decline, indicating no phage-mediated bacterial lysis during the non-replicating state in PBS **(Supp Fig 3)**. To further distinguish whether phage-mediated killing of bacteria is occurring during the infection or during the revival of bacteria on 7H10 plates, we carried out a resazurin metabolic assay of phage exposed and non-phage exposed bacteria in PBS and found no difference in metabolic activity between the two conditions, suggesting that the phage did not kill the bacteria in the non-replicating state and instead phage-mediated killing happened during the revival of bacteria on 7H10 plates **(Fig 2B)**. Interestingly, we also observed a time-dependent decay of phage efficacy with no significant difference observed between phage-exposed and non-exposed groups post 25 days of phage-exposure (**Fig 2A**). To evaluate whether this reduced bacterial revival on 7H10 agar is because of the formation of Viable But Non-Culturable bacteria (VBNCs)^46^ in phage-exposed populations, we carried out a Most Probable Number (MPN) assay^47^ and found no differential increase in total viable bacteria in the two groups **(Supp Fig 4)**. We next checked the efficacy of antibiotics against non-replicating *M. smegmatis*. We used a combination of three antibiotics above their MICs for *M. smegmatis* - Rifampicin (30 µg/mL, 1.2x MIC), Isoniazid (20 µg/mL, 1.3x MIC), and Ethambutol (1 µg/mL, 5x MIC) - three of the four front-line drugs for the treatment of Tuberculosis. We found that the combination of the three antibiotics was ineffective in killing non-replicating bacteria **(Supp Fig 5)**, demonstrating the higher potential of phages to persist with non-replicating bacteria and lyse them upon revival.

**Figure 3.**
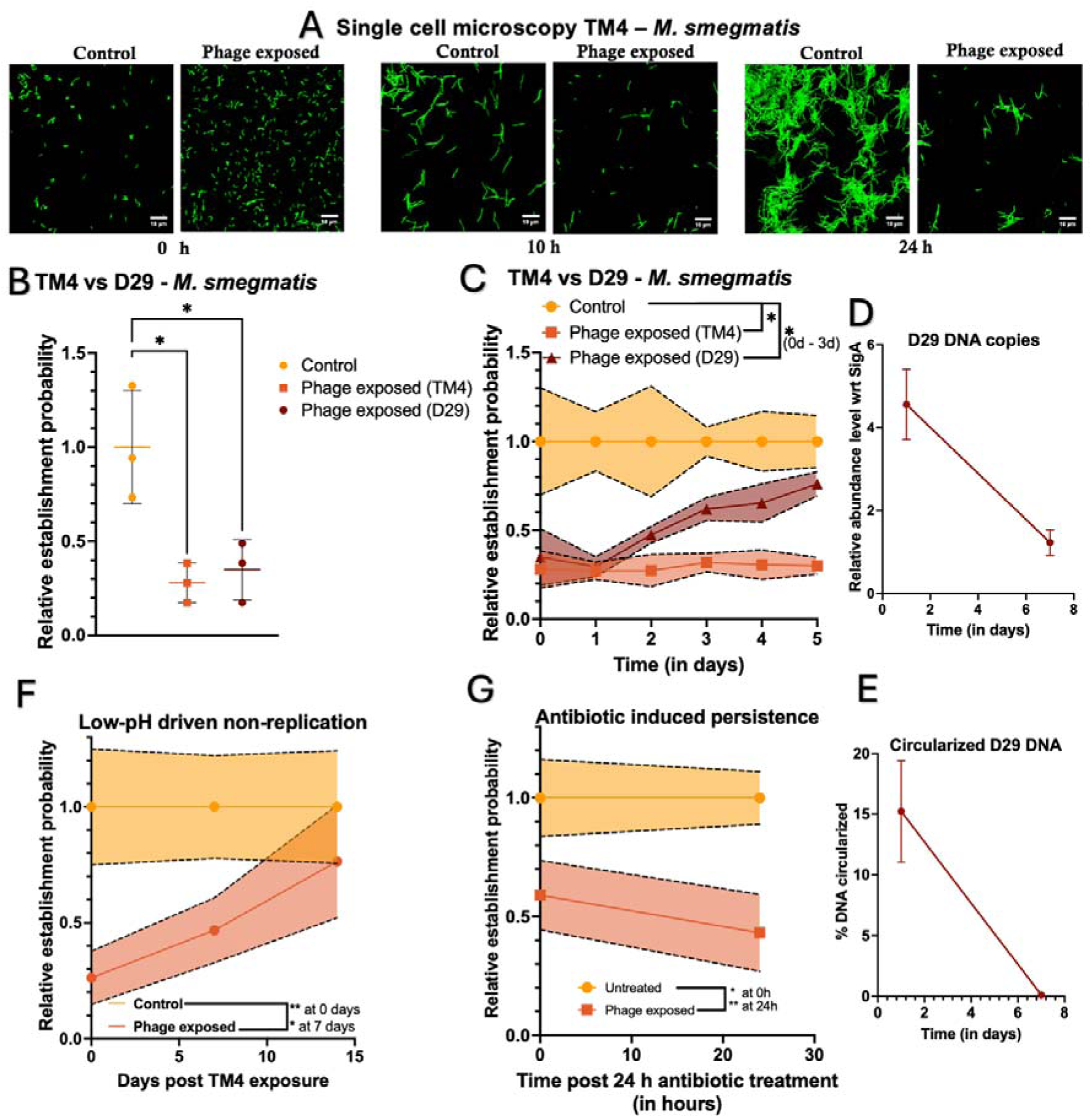
Single-cell growth analysis of phage-exposed bacteria. **A)** Single cell timelapse microscopy of non-replicating M. smegmatis exposed to TM4 bacteriophage for 3 hours prior to growth media supplementation (Scale bar: 10 µm) (n = 4). **B)** TM4 vs D29 (10:1 MOI) phage infectivity and efficacy towards non-replicating M. smegmatis (n=3). **C)** TM4 vs D29 (10:1 MOI) pseudolysogeny window in non-replicating M. smegmatis (n=3). **D)** Quantitative PCR of total genomic DNA extracted from phage-exposed M. smegmatis cultures to detect and quantify D29 phage DNA (n=3) and **E)** D29 phage DNA stability quantified through circularization of phage DNA (n=3). **F)** TM4 (10:1 MOI) phage efficacy against low-pH (pH 4.5) driven non-replicating M. smegmatis (n=3). **G)** TM4 (10:1 MOI) phage efficacy against antibiotics (Rifampicin, Isoniazid and Ethambutol at 2X MIC) induced persistence in M. smegmatis (n=3). For B, unpaired t-test was used to determine the statistical significance. For C, F, and G, 2-way ANOVA with Sidak’s multiple comparisons test was used to determine statistical significance. (n=3). (*p ≤0.05, **p ≤0.01, ***p ≤0.001, ****p ≤0.0001).

**Figure 4.**
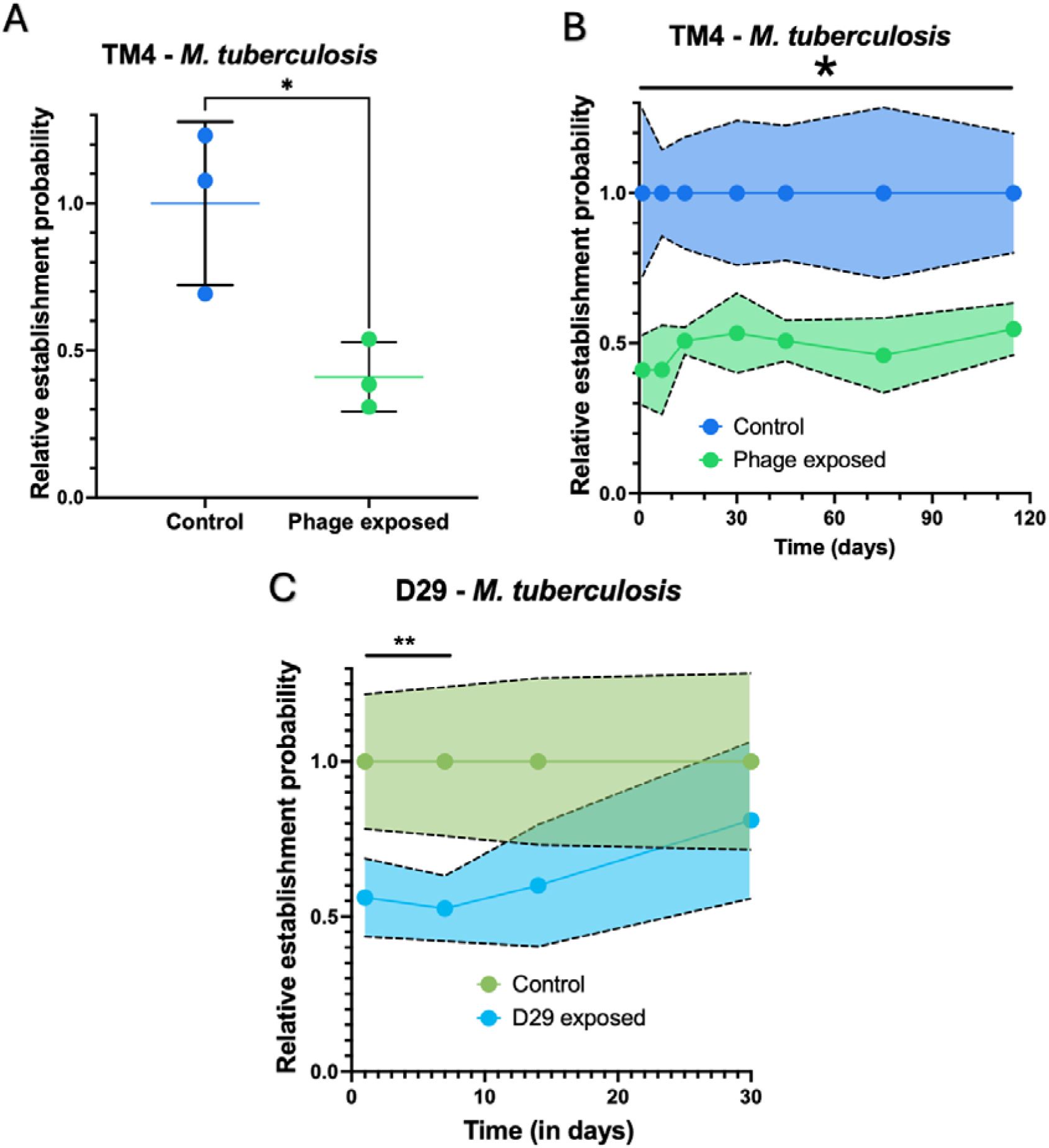
Efficacy of phage pseudolysogeny across different species of host and phage. **A)** TM4 phage infectivity and efficacy towards non-replicating M. tuberculosis (H37Rv) quantified through single-cell growth assay (n=3) **B)** TM4 pseudolysogeny window in non-replicating M. tuberculosis quantified through single-cell growth assay (n=3) **C)** D29 pseudolysogeny window in non-replicating M. tuberculosis quantified through single-cell growth assay (n=3). For A, C unpaired t-test was used to determine the statistical significance. For B, two-way ANOVA with Sidak’s multiple comparisons test was used to estimate the statistical significance (*p ≤0.05, **p ≤0.01, ***p ≤0.001, ****p ≤0.0001).

**Figure 5.**
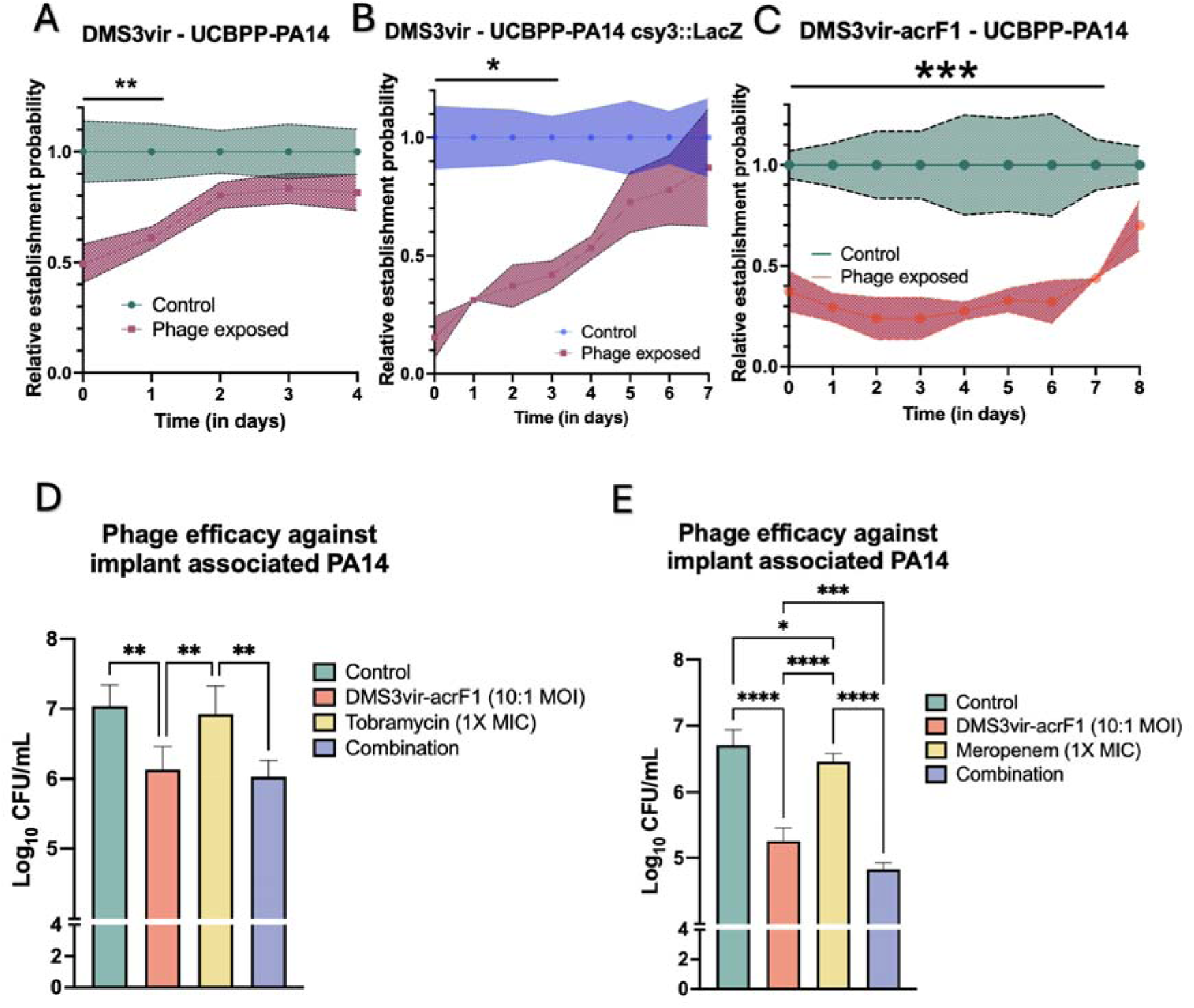
CRISPR activity in regulating phage pseudolysogeny in nutrient stressed host. **A)** CRISPR susceptible DMS3vir phage pseudolysogeny window in nutrient stressed CRISPR-active P. aeruginosa quantified through single-cell growth assay (n =3) **B)** CRISPR susceptible DMS3vir phage pseudolysogeny window in nutrient stressed CRISPR-inactive P. aeruginosa quantified through single-cell growth assay (n =3) **C)** CRISPR resistant DMS3vir-AcrF1 phage infectivity and efficacy towards nutrient stressed CRISPR-active P. aeruginosa quantified through single-cell growth assay (n =3) **D)** Efficacy of Tobramycin (1X MIC) and DMS3vir-AcrF1 (10:1 MOI) individually and in combination on nutrient stressed P. aeruginosa on implant surface (n=3) **E)** Efficacy of Meropenem (1X MIC) and DMS3vir-AcrF1 (10:1 MOI) individually and in combination on nutrient stressed P. aeruginosa on implant surface (n = 3). For A, B and C, two-way ANOVA with Sidak’s multiple comparisons test was used to assess statistical significance. For D, E unpaired t-test was used to determine the statistical significance (*p ≤0.05, **p ≤0.01, ***p ≤0.001, ****p ≤0.0001).

### 4. Single-cell analysis revealed phage-mediated lysis in *M. smegmatis* occurred upon bacterial resuscitation

A limitation of ensemble-based CFU enumeration of bacterial survival is that other confounding factors, such as quorum sensing and phage diffusion across the agar plate, could result in an inaccurate estimation of phage efficacy. To address these limitations, we analysed the proliferation of single *M. smegmatis* cells using principles of Poisson distribution^48^. We infected 48 h PBS-starved *M. smegmatis* cultures with TM4 bacteriophage at 10:1 MOI for 3 h and removed extracellular phage. At regular intervals, we seeded approximately 1 bacterial cell/well in a 96-well plate. This was achieved by diluting the starved cultures to around 100 cells/mL. At these high dilutions, the distribution of cells is governed by the Poisson distribution. A well with seeded bacterium could fail to turn turbid either due to the absence of any bacterium in the inoculum or due to the bacterium’s inability to produce a surviving lineage. Wells receiving no cells were accounted for by normalizing the phage-exposed group to the non-exposed group. The plates were incubated at 37 °C for 7 days, after which the number of wells showing growth was counted. As the phages after FAS treatment were 3-4-log order lower compared to bacteria, we do not expect any free extracellular phage to interfere in this assay after dilutions to 1 bacterium per well. Similar to the bulk kinetics, we observed ∼60-70% reduction in the ability of a single non-replicating *M. smegmatis* cell to proliferate when exposed to phages **(Fig 2C)**. Furthermore, we measured the pseudolysogeny window of TM4 bacteriophage in non-replicating *M. smegmatis* to be around 14 days after which the phage effectiveness reduced with time **(Fig 2D)**, in agreement with the bulk CFU results **(Fig 2A)**. Phage detection through PFU assay of 96-well plate samples showed a higher probability of phage presence initially up to 14 days with gradual decline upto day 35 **(Fig 2E)**. We also validated these observations with single-cell microscopy analysis of agarose pad-trapped *M. smegmatis.* Microscopy analysis revealed a similar trend in the inability of phage-exposed *M. smegmatis* to revive from a non-replicating state and the prolonged pseudolysogeny of TM4 bacteriophage in non-replicating *M. smegmatis* **(Fig 3A, Supp Fig 6)**. To understand whether the bacterial growth (∼40%) observed in the phage-exposed samples at early time points was because of a fraction of the bacteria not being infected with phage **(Fig 2D)**, we fluorescently tagged bacteriophage TM4 DNA with SYBR Gold^49^ prior to bacterial exposure. We then performed Flow Assisted Cell Sorting (FACS) and collected SYBR-Gold positive (bacteriophage-infected) cells and analysed single-cell proliferation. Interestingly, we observed a significant difference in relative establishment probability (REP) (REP reduced to ∼0.1 compared to 0.4) in phage-exposed and flow-sorted vs non-sorted bacteria **(Supp Fig 7),** suggesting a significant proportion of the bacterial growth observed at the early point could be attributed to the inability of phage to infect all bacterial cells or initiate its gene expression.

**Figure 6.**
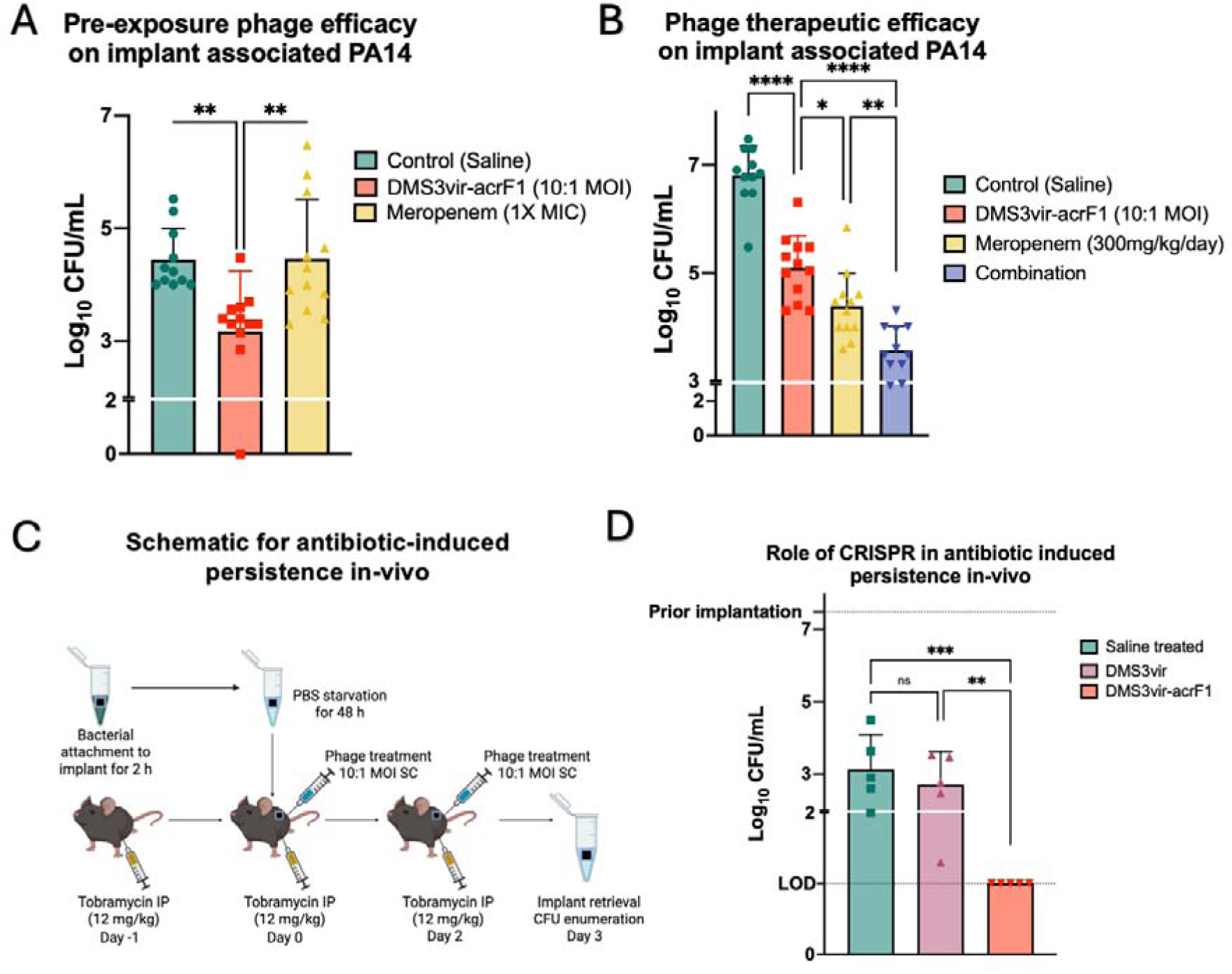
Phage efficacy against nutrient-stressed implant associated P. aeruginosa in vivo. **A)** In vivo pre-infection efficacy of Pseudomonas phage DMS3vir-AcrF1 in reducing implant associated bacterial counts in Balb/c mouse model (n=10, data pooled from two independent experiments) **B)** In vivo treatment of implant associated Pseudomonas infections in C57BL/6 mice (n=10, data pooled from two independent experiments) **C)** Schematic representing in vivo antibiotic driven persistence of implant associated P. aeruginosa **D)** In vivo treatment of antibiotic driven persistent implant-associated P. aeruginosa infections in C57BL/6 mice with CRISPR-resistant (DMS3vir-acrF1) and CRISPR-susceptible (DMS3vir) bacteriophages (representative data) (n=5). Prior implantation dotted line indicates the CFU of bacteria on the implant prior to implantation in antibiotic injected mice. LOD dotted line (10 bacteria) is the limit of detection through CFU assay. Ordinary one-way ANOVA was used to estimate the statistical significance. (*p ≤0.05, **p ≤0.01, ***p ≤0.001, ****p ≤0.0001). Each experiment was repeated twice.

To check whether pseudolysogeny and its prolonged maintenance are phage-specific phenomena, we used another lytic bacteriophage, D29, to check for infectivity and efficacy against non-replicating *M. smegmatis*. Phage infection efficacy of D29 in log-phase *M. smegmatis* is equivalent to TM4, as expected **(Supp Fig 8)**. D29 was able to infect non-replicating *M. smegmatis***(Fig 3B)** with slightly lower efficacy than TM4 at around 5 D29 DNA copies per bacterial DNA copy **(Fig 3D).** Its pseudolysogeny window was much shorter than that of TM4, with DNA copies, circularization and bacterial lysis reducing beyond day 1 **(Fig 3C-E)** suggesting that although the phenomenon of pseudolysogeny might exist in several phages, the phage DNA stability is bacteriophage specific.

### 5. Phage pseudolysogeny remained efficacious in low-pH induced non-replication as well as in antibiotic-driven persistence

Intracellular pathogens such as *Staphylococcus aureus* and *M. tuberculosis* are able to survive in a low-pH environment and antibiotic challenge by adopting a non-replicating state^50,51^. To understand whether the non-replication state induced by low pH would affect the ability of bacteriophage to infect and persist, we employed a low-pH model for non-replicating *M. smegmatis*. The growth of bacteria was tested at different pH and at pH 4.5, *M. smegmatis* entered a state of non-replication as shown by the clock plasmid **(Supp Fig 9A)**. We proceeded to test phage infectivity and efficacy against non-replicating *M. smegmatis* by infecting with TM4 bacteriophage at 10:1 MOI at pH 4.5 and quantifying through single-cell proliferation assay. We found that bacteriophage TM4 remains infective towards *M. smegmatis* at pH 4.5 **(Supp Fig 9B)** and is successful in lysing the bacterial cells upon reactivation for upto 7 days (**Fig 3F**), suggesting that pseudolysogeny is active under diverse host environmental conditions that induce non-replication.

Antibiotic treatment is known to enrich for metabolically dormant (persister) cells within a bacterial population^52^. To assess whether bacteriophages can infect and eliminate such persisters, *M. smegmatis* cultures were grown in nutrient-rich media until mid-log phase. Cultures were then treated with rifampicin–isoniazid–ethambutol (2x MIC) for 24 h, after which CFUs significantly dropped and plateaued **(Supp Fig 9C)**, indicating persister enrichment. Subsequent exposure to TM4 (MOI = 10) for 3 h led to a pronounced decline in CFUs **(Fig 3G)**, confirming phage infectivity and killing of antibiotic-tolerant cells. Optical densities (OD) of cultures remained unchanged, suggesting killing occurred during revival on antibiotic free media plates.

### 6. Phages were effective against virulent non-replicating *M. tuberculosis*

To determine whether this phenomenon of pseudolysogeny is species-specific to *M. smegmatis*, we assessed the infectivity of TM4 and D29 bacteriophages against non-replicating *M. tuberculosis*. Replication rates of *M. tuberculosis* are known to rapidly drop over a 96 h period of nutrient starvation and the bacteria enter a non-replication state subsequently^53^. Log-phase secondary cultures of *M. tuberculosis* were resuspended in PBS and incubated at 37 °C, 180 RPM for one week. The cultures were then exposed to either TM4 or D29 bacteriophage for 3 h, followed by AFS treatment. Periodically, cultures were aliquoted and growth was quantified using single-cell growth method. Bacteriophage TM4 remained infective against non-replicating *M. tuberculosis* **(Fig 4A)** and continued to be effective for over 120 days **(Fig 4B)** whereas, similar to *M. smegmatis*, D29 lost its efficacy in under a week **(Fig 4C)**.

### 7. Pseudolyosogeny demonstrated in *Pseudomonas aeruginosa* and CRISPR defence played an active role in phage neutralization

Although bacteria have evolved diverse anti-phage defence strategies, bacteriostatic conditions have been shown to preferentially favour the development of CRISPR-Cas adaptive immunity over resistance mediated by surface modifications^35^. Furthermore, slow-replicating phages may be more vulnerable to CRISPR-Cas interference, due to increased protospacer acquisition^54^. Therefore, to understand the role of CRISPR-Cas adaptive immunity in bacteria against phage pseudolysogeny and to determine the pseudolysogeny stability in other bacterial genera, we studied the phage infectivity of nutrient-starved *Pseudomonas aeruginosa* (UCBPP-PA14) which harbours an active CRISPR-Cas system unlike *M. smegmatis* which lacks a CRISPR-Cas system^55^. First, we checked the replication levels of 48-hour PBS-starved *P. aeruginosa* prior to checking phage infectivity of CRISPR-Cas3 Type I-F susceptible DMS3vir against nutrient-stressed *P. aeruginosa*, against the wild-type strain (UCBPP-PA14) and the CRISPR inactive mutant (UCBPP-PA14 csy3::LacZ)^56,57^. We found that even after 48 h of stringent nutrient starvation, unlike *M. smegmatis*, *P. aeruginosa* was able to replicate, in contrast to previous reports describing deep dormancy in 48 h stationary-phase cultures^41^ **(Supp Fig 10A)**. However, single-cell microscopy-based quantification revealed that less than ∼30% of cells were actively dividing **(Supp Fig 10B)**. Increasing the nutrient starvation period for up to 12 days did not improve the fraction of non-replicating bacteria. Given that the majority of the population had entered a non-replicative state 48 h post nutrient-starvation, we next examined phage infectivity towards these bacteria. The phage DMS3vir was infective towards both wild-type and CRISPR-inactive strains of PA14 **(Supp Fig 11A, B)**. However, the phage DNA was quickly lost (in 1-2 days) in the wild-type strain containing an active CRISPR-Cas system **(Fig 5A)**. The role of CRISPR in phage efficacy loss was further confirmed by RT-PCR to check for protospacer sequence acquisition during the loss of phage efficacy, which revealed the activity of the PA14 CRISPR-Cas system even under stringent starvation conditions **(Supp Fig 12)**. The pseudolysogeny window in the CRISPR-inactive strain was extended for nearly a week **(Fig 5B)**. Although CRISPR inactivation extends the pseudolysogeny phase of CRISPR-susceptible phages, this approach is difficult to implement in clinical settings. Alternatively, bacteriophage modifications can be employed to confer resistance to CRISPR interference. To evaluate whether such phage modifications can prolong the pseudolysogeny phase in CRISPR-active, nutrient-starved bacteria, we utilized the CRISPR-resistant bacteriophage DMS3vir-AcrF1^58^ to investigate the pseudolysogeny period in CRISPR-active, nutrient stressed *Pseudomonas aeruginosa*. The CRISPR-resistant bacteriophage DMS3vir-acrF1 infected nutrient stressed *P. aeruginosa* strain UCBPP-PA14 **(Supp Fig 11C)**, resulting in a protracted pseudolysogeny period of upto 7 days **(Fig 5C)** – demonstrating the ability of CRISPR-resistant bacteriophages in circumventing CRISPR activity even under stringent nutrient starvation conditions. To further confirm this increase in pseudolysogeny window was because of a lack of CRISPR response, we checked for protospacer sequence acquisition after 24 h **(Supp Fig 12)** which showed no protospacer acquisition in DMS3vir-AcrF1 exposed bacteria.

*P. aeruginosa* readily transitions into a persistent state following antibiotic exposure in planktonic cultures^59^. To assess phage efficacy against these antibiotic-induced persisters, mid-log–phase cultures grown in nutrient-containing media were treated with 10× MIC meropenem and 5× MIC tobramycin for 24 h. CFU counts subsequently plateaued **(Supp Fig. 13A)**, indicating enrichment of a persistent subpopulation. When these cultures were exposed to DMS3vir-acrF1 phages (MOI = 10) for 3 h, CFUs declined markedly (**Supp Fig. 13B**), demonstrating phage infectivity and killing of antibiotic-driven persistent cells on revival. Phage PFUs gradually declined over multiple days indicating no phage replication on antibiotic-driven persistent *P. aeruginosa* **(Supp Fig. 13C)**. Notably, optical densities remained stable, suggesting that killing occurred upon revival. This CFU reduction persisted over 24 h (**Supp Fig. 13B**).

### 8. Phages infected implant-associated bacteria and caused lysis upon revival

Implant-associated *P. aeruginosa* infections have been reported to have relapse rates as high as 30%^60^. To evaluate the clinical relevance of phage pseudolysogeny towards the prevention of infection relapses, an implant-associated *P. aeruginosa* infection model was employed. Following incubation of *P. aeruginosa* (UCBPP-PA14) with stainless steel (SS316L) for 2 h, the implant-associated bacteria were starved for 48 h **(Supp Fig 14A).** Bacterial adhesion to abiotic surfaces is improved by the presence of ions-such as Mg^2+61^ and in order to facilitate high bacterial adhesion to the implants, starvation for *in vitro* experiments was carried out in relatively ion-rich M9 salts (no carbon provided). To simulate the clearance of antibiotics and phage over time observed in clinics, we incorporated a latency step where the implant was again placed in M9 salts without antibacterials for another 24 h **(Supp Fig 14A)**. Upon enumeration of the CFUs of implant-associated populations on LB-agar growth medium, we observed that tobramycin at 1×MIC did not result in reduction in CFU while meropenem resulted in a modest reduction in bacterial load compared to untreated controls **(Fig 5D, E)**. Treatment with the CRISPR-resistant phage DMS3vir-AcrF1 (10:1 MOI) alone resulted in a reduction of bacterial counts by approximately one log.

When phages were combined with meropenem, we observed an additive effect in these conditions **(Fig 5E)**. DM3vir-acrF1 was also able to successfully infect and lyse non-replicating implant-associated multidrug-resistant clinical isolate of *P. aeruginosa* BEI 12368 resistant to imipenem, meropenem, kanamycin, neomycin, butirosin, seldomycin, chloramphenicol, fosfomycin and beta-lactams **(Supp Fig 15)**. Overall, phage monotherapy showed greater efficacy than antibiotic monotherapy, and combination treatment was only marginally beneficial in the case of meropenem.

### 9. In vivo relevance of pseudolysogeny in an implant infection model

Implant-associated *P. aeruginosa* infections are among the most difficult to eradicate due to biofilm formation, metabolic heterogeneity, and intrinsic antibiotic resistance, often necessitating implant removal for cure^62,63^. In order to understand if this pseudolysogenic efficacy of DMS3vir-acrF1 can be translated *in vivo*, we developed a mouse model of *Pseudomonas aeruginosa* (UCBPP-PA14) implant-associated infection. The effect of pseudolysogeny was studied in this model through a series of mouse experiments. Resuscitation of non-replicating bacteria *in vitro* typically occurs in uniform, nutrient-rich conditions, whereas *in vivo* reactivation is constrained by heterogeneous microenvironments characterized by nutrient gradients, immune-mediated stress, and tissue-specific factors. First, to evaluate whether phage pseudolysogenized non-replicating bacteria would be susceptible to phage lysis *in vivo*, we exposed bacteria to DMS3vir-acrF1 at an MOI of 10:1 prior to implant attachment. Subsequently, infected implants were subcutaneously placed in mice. The number of implant-associated bacteria was significantly reduced two days post-implantation **(Fig. 6A, Supp Fig 14B)**. Treatment of non-replicating bacteria with 1× MIC meropenem prior to implant attachment did not reduce bacterial counts *in vivo*, further highlighting the inefficacy of antibiotics against non-replicating bacteria **(Fig. 6A)**. Next, to evaluate the phage therapeutic potential, we tested the efficacy of phage treatment post-implantation in mice. Bacteria were allowed to attach onto the implants for 2 h, following which the implant-associated bacteria were nutrient-starved in PBS for 48 h. The implants were then placed subcutaneously in mice, followed by the treatment with DMS3vir-acrF1 or meropenem (300 mg/kg/day) subcutaneously near the site of implantation. Both phage and antibiotic treatment, either individually or in combination rescued the mice from succumbing to infection-related death and reduced the bacterial counts associated with the implant **(Fig 6B)**. The combination treatment showed a significant synergistic effect compared to the individual treatments, highlighting the ability of phages to be used as a cotreatment strategy in treating implant-associated bacterial infections **(Fig 6B)**. Together, these results suggest that phage pseudolysogeny remains active during bacterial resuscitation from a non-replicating state *in vivo* and phage therapy, especially in combination with antibiotics could be a viable strategy for targeting implant-associated bacteria.

### 10. Phage efficacy in an antibiotic-driven implant associated persistent infection model

Antibiotic-induced persistence is a major contributor to infection relapse^64,65^. Such persister cells exhibit markedly reduced metabolic activity and replication, rendering antibiotics ineffective. To assess phage efficacy against antibiotic-induced persisters, we established an implant-associated *P. aeruginosa* infection model in mice. Log phase *P. aeruginosa* were allowed to adhere to SS316L stainless steel implants for 2 h, washed, and resuspended in PBS for 48 h prior to implantation. C57BL/6 mice received intraperitoneal tobramycin (12 mg/kg/day) starting one day before implantation surgery and continuing the injection regimen for the rest of the days to maintain antibiotic pressure. The CFU counts also demonstrated that under this daily antibiotic pressure, the bacterial numbers reduced by several log-folds compared to previous animal experiment and then remained stable *in vivo* **(Supp Fig. 16)**. To check for the relevance of the CRISPR-Cas system *in vivo* in phage therapy, the implants were placed into the mice subcutaneously, and the mice were administered saline, CRISPR susceptible DMS3vir (MOI 10/day) or CRISPR resistant DMS3vir-acrF1 phage (MOI 10/day) subcutaneously, near the implantation site. Implants were retrieved 2 days post-surgery, and adherent bacteria were quantified. Remarkably, implants from mice receiving CRISPR-resistant phage harboured no bacteria (at LOD) while saline-treated and CRISPR-susceptible phage-treated mice implants harboured bacteria in equivalent amounts **(Fig 6C)**. From a clinical perspective, this highlights the potential of optimized phage design to prevent chronic and persistent implant-associated bacterial infections and reduce the need for prolonged antibiotic therapy or surgical revision, especially in cases of infection relapse.

## DISCUSSION

Recurrent bacterial infections remain a major clinical challenge, driven in part by antibiotic-persister cells. Emerging evidence suggests that persistent bacteria play a major role in many recurrent infections. Invasive non-typhoidal Salmonella infections, for example, relapse in 78% of HIV patients due to persistence rather than reinfection^65^ with similar trends observed in UTIs^66^, and Pseudomonas infections^67^. *Mycobacterium tuberculosis* infections are known to persist even after the completion of antibiotic regimens, leading to relapse in approximately 10% of patients over their lifetime^7,68^. *Pseudomonas aeruginosa* in nosocomial settings shows relapse rates of 12–18%^6^. These persistent populations promote the evolution of antibiotic resistance^69,70^, further highlighting the need to study and design therapies against bacterial persistence. Bacteria survive stress through diverse mechanisms, including (p)ppGpp-mediated replication arrest^71^; the oxidative stress response (SoxS-SoxR)^72^,contact-dependent growth inhibition^73^, and ATP depletion^74^. These adaptations enable dormancy under host, antibiotic, or nutrient stress, allowing survival at otherwise lethal antibiotic levels without heritable resistance. The resilience of such tolerant populations has intensified interest in alternative therapeutics, notably lytic bacteriophages with potent, species-specific bactericidal activity^75^. Upon binding to bacterial surface receptors, lytic phages inject their genetic material and hijack the host’s replication machinery, ultimately leading to host cell lysis^20^. However, the low metabolic state of bacteria can prevent phage replication in non-replicating bacteria, resulting in a state of pseudolysogeny^20,76^. In a couple of studies by Miller and Ripp^29,76^, it has been observed that *Pseudomonas* lytic bacteriophage UT1 is able to undergo pseudolysogeny when exposed to nutrient-starved bacteria and is able to resume replication and produce phage virions upon nutrient supply post 24h. Bryan et al. observed that T4 bacteriophage is able to infect stationary phase *E. coli* and enter “hibernation” mode until nutrient supply is restored^21^. We observed a similar phenomenon with Mycobacteriophage TM4 and D29 **(Fig 2A, 2C and 3A)** and *Pseudomonas* phage DMS3vir and DMS3vir-acrF1 **(Supp Fig 12)**, wherein the phages retained infectivity but failed to replicate within non-replicating host cells. However, no previous studies examined how long this pseudolysogeny can persist for in a stressed bacterial cell, the role of bacterial defence systems under these conditions and its therapeutic relevance in pre-clinical models.

In our study, we found phage reactivation post nutrient supplementation occurred only upto a specific time; after which, phage efficacy was lost **(Fig 2D**, **Fig 3B**, **Fig 4D**, **Fig 5A-B)**. This decline may result from either the gradual degradation or loss of phage DNA within the non-replicating bacteria during extended starvation or due to an active defence system eliminating the foreign DNA. Consistent with this, we observe a time-dependent decrease in phage DNA copy number in starved cells **(Fig 1C-D**, **Fig 4A)**. We further investigated the pseudolysogeny windows of mycobacteriophages TM4 and D29 in two mycobacterial species: *Mycobacterium smegmatis* mc²155 and *Mycobacterium tuberculosis* H37Rv. TM4 exhibited a pseudolysogeny window of 14 days in *M. smegmatis* **(Fig 2D)**; however, in *M. tuberculosis*, this window was markedly prolonged—persisting for up to 120 days of bacterial starvation, at which point the experiment was terminated **(Fig 4C)**. In contrast, D29 showed a much shorter pseudolysogeny window of only a few days in both species **(Fig 3B, Supp Fig 10)**. Correspondingly, D29 phage copy numbers were significantly lower, suggesting reduced stability and infectivity of D29 in non-replicating *Mycobacterium spp* **(Fig 4A)**. This difference in infectivity of TM4 and D29 bacteriophages towards non-replicating bacteria could be due to the previously reported ability of TM4 to efficiently infect stationary phase Mycobacterial cells through the mt3 motif in the tail measure protein^77^, whereas D29 is unable to infect stationary phase cells through mt3 motif. Our findings further confirm the previous reports that D29 bacteriophage has poor infectivity towards stressed *Mycobacterium*^78^.

While nutrient starvation and the formation of non-replicating persister cells are well-studied bacterial stress responses, other stressors, such as low pH and antibiotic exposure, also contribute significantly to bacterial persistence and disease relapse^51,79,80^. To investigate whether pseudolysogeny enables phage survival under such diverse conditions, we evaluated the efficacy of phages against low pH- and antibiotic-induced non-replicating *Mycobacterium smegmatis*. In both scenarios, phages remained effective **(Fig 3C, 3D)**, supporting the hypothesis that pseudolysogeny serves as one of the key mechanisms by which phages persist in diverse microenvironments^29^. To explore the role of host immunity in limiting pseudolysogeny, we investigated the contribution of CRISPR-Cas systems. Approximately 40% of bacterial genomes encode CRISPR-Cas loci, which constitute a major form of adaptive immunity against phages^81^. While previous studies have shown that CRISPR-Cas systems remain active even under bacteriostatic conditions^35^, their impact on phage pseudolysogeny remains poorly understood. Prior reports with *Pseudomonas aeruginosa* and DMS3vir show that bacteria grown in nutrient-limited media or in the presence of bacteriostatic antibiotics show evolution towards a CRISPR-driven phage resistance^35^. In agreement with these reports, we find that the CRISPR-Cas system in *Pseudomonas aeruginosa* effectively suppresses pseudolysogeny in nutrient-starved conditions **(Fig 5A)**. However, phages expressing anti-CRISPR proteins, such as DMS3-AcrF1, were able to overcome this defence and maintain a pseudolysogenic state **(Fig 5B, C)**. To our knowledge, this is the first demonstration of anti-CRISPR phages sustaining pseudolysogeny in non-replicating bacteria, thereby expanding the therapeutic potential of engineered phages in treating persistent infections.

To further assess the therapeutic potential of phage pseudolysogeny, we developed an implant-associated, non-replicating Pseudomonas mouse infection model. In this system, Pseudomonas aeruginosa cells were allowed to adhere to an implant surface, which was then surgically implanted into mice. To ensure that implant-associated bacteria in our animal model remained stressed and predominantly non-replicating, mice received continuous tobramycin treatment throughout the study, which resulted in stable CFU counts **(Supp Fig. 18)**. Our *in vivo* results show that phage plays a significant role in reducing antibiotic-tolerant bacterial populations **(Fig 6C, D)**. Remarkably, we observed high efficacy of CRISPR-resistant phage in controlling implant-associated bacteria **(Fig 6C, D)** compared to a CRISPR-susceptible phage.

Although previous studies show the efficacy of anti-CRISPR phages in reducing bacterial burden in mice^82^ and a role of CRISPR defence in reducing phage efficacy *in vivo* against gut bacteria^83^, to our knowledge, this is the first study to evaluate the therapeutic impact of phage pseudolysogeny lifecycle as well as the role of CRISPR immunity in phage therapy against persistent bacteria. These findings support a new direction in phage therapy design—incorporating pseudolysogeny-capable CRISPR-resistant phages into therapeutic cocktails to more effectively target and eliminate persistent infections.

This study has a few limitations. First, while this work demonstrated the ability of CRISPR-resistant phages to overcome CRISPR-mediated immunity, bacteria encode multiple additional defence systems whose contributions to phage pseudolysogeny and therapeutic efficacy were not explored here. Second, although differential phage stability was observed in non-replicating hosts, the precise molecular mechanisms governing phage genome persistence, stabilization, or degradation remain unresolved. Third, it remains unknown whether phage pseudolysogeny can be directly modulated using small-molecule/drug interventions; at present, modulation was only observed through evolutionary advantages conferred to phages, such as resistance to CRISPR-mediated targeting. Finally, phage carrier-states and pseudolysogeny are sometimes speculated to promote horizontal gene transfer (HGT) in bacteria due to the prolonged presence of the phage genome within the host. Although evidence exists for homologous recombination between phage and bacterial genomes^84^, no direct evidence exists for HGT mediated by carrier-states or phage pseudolysogeny and further research is necessary to understand HGT mediated by these alternate phage lifestyles, especially in the context of lytic phages. Addressing these questions in future studies will be essential for refining phage selection strategies and for rationally designing therapies that exploit phage–host interactions under persistent infection states.

## CONCLUSION

We demonstrated that lytic bacteriophages can infect and persist within persistent bacteria via pseudolysogeny, ultimately lysing bacteria upon reactivation. The duration of this pseudolysogenic state is both phage- and host-dependent, influenced by phage DNA stability and host replication rates, calling for a rational phage therapy design especially for the treatment of persistent bacterial infections. Moreover, we report that host immune systems like CRISPR-Cas can restrict pseudolysogeny even under non-replicating state, but anti-CRISPR strategies offer a viable workaround. We believe these findings lay a systematic groundwork for leveraging phage biology and its interaction with non-replicating bacteria as a novel therapeutic approach to target and improve the treatment of persistent, relapse-prone bacterial infections.

## ACKNOWLEDGEMENTS

We thank Dr. Sujoy K. Das (Bose Institute, Kolkata) and Dr. Graham Hatfull (University of Pittsburgh) for providing us with mycobacteriophages D29 and TM4 respectively. We thank Dr. William R. Jacobs (Albert Einstein College of Medicine) for providing us with fluorophage (φ^2^GFP10). We thank Dr. Edze Westra (University of Exeter, United Kingdom) for kindly providing us with the CRISPR inactive *P. aeruginosa* and CRISPR-resistant *Pseudomonas* phages used in this study. We also thank Dr. Narendra M. Dixit and Dr. Amit Singh for providing valuable feedback on the manuscript. The following reagent was obtained through BEI Resources, NIAID, NIH: Pseudomonas aeruginosa, Strain MRSN 12368, NR-51574. This strain is part of the Pseudomonas aeruginosa Diversity Panel provided by the Multidrug-Resistant Organism Repository and Surveillance Network (MRSN) at the Walter Reed Army Institute of Research (WRAIR). We thank the Department of Bioengineering, IISc, Flow cytometry facility, IISc, Centre for Infectious Disease Research, IISc for instrument and facility access.

Funding: Bill & Melinda Gates Foundation [OPP1210498] and Indian Council for Medical Research [IIRPSG-2024-01-02611] to Rachit Agarwal, Prime Minister’s Research Fellowship [PMRF ID: 0201550] to Yeswanth Chakravarthy Kalapala, Kishore Vaigyanik Protsahan Yojana Fellowship [KVOY ID: X46010020] to Aditya Kamath Ammembal, Department of Biotechnology – Junior Research Fellowship [DBTHRDPMU/JRF/BET-24/I/2024-25/214] to Saksham Jain, Prime Minister’s Research Fellowship [PMRF ID: 0202578] to Nisha Sanjay Barge.

## CONTRIBUTIONS

R.A. and Y.C.K. conceptualized and developed the study methodology. Y.C.K. performed experiments with *M. smegmatis*, *M. tuberculosis* and *P. aeruginosa.* A.K.A. performed and optimized implanted associated *P. aeruginosa in vitro* experiments. Y.C.K. and A.K.A. performed *in vivo* studies with the assistance of S.J. and N.S.B. S.J. analysed the RNA expression data.

## COMPETING INTERESTS

The authors declare no competing interests.

## METHODS

### Bacterial Cell Culture and Maintenance

Primary *M. smegmatis* and *M. tuberculosis* were grown in Middlebrook 7H9 broth (Sigma, St Louis, MO, United States) supplemented with Glycerol, ADC and Tween-80 (0.1% v/v). Secondary cultures of *M. smegmatis* and *M. tuberculosis* were grown in 7H9 broth supplemented as above without Tween-80. *Pseudomonas aeruginosa* cultures were grown in LB media (Himedia, Maharashtra, India). For starvation, 1X PBS was used for *M. smegmatis* and *M. tuberculosis* and 1X PBS or M9 salts were used for *P. aeruginosa* (Himedia, Maharashtra, India).

### Bacterial nutrient starvation and non-replication enumeration

Log phase secondary cultures of *M. smegmatis* containing clock plasmid pBP10^36^ were centrifuged at 5000 g for 5 mins at room temperature and the bacterial pellet is washed twice with PBS before resuspending in PBS solution. The cultures were stored at 37 °C with 180 RPM shaking and at regular intervals, aliquots were drawn and serially diluted to enumerate the Colony Forming Units (CFUs) on 7H10 plates or on Kanamycin (30 μg/mL) containing 7H10 plates. Percentage of bacteria with plasmid was calculated from both the CFU counts using the following formula:

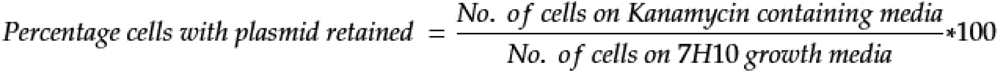

### Phage infection check and optimization

□^2^GFP10 bacteriophage^39^ encoding GFP gene was employed to study the ability of the phage to infect non-replicating bacteria. Briefly, 48 h PBS starved *M. smegmatis* cultures expressing RFP were infected with 10:1 MOI of □^2^GFP10 at 37 °C for various timepoints. At each point, the cultures were centrifuged at 5000 g for 5 min and the pellet was washed with MilliQ H_2_0 before resuspending the culture in 7 mM Ammonium Ferric Sulphate (AFS) to rid any extracellular phage^45^. Post AFS treatment, the culture was washed once with milliQ and once with PBS and resuspended in PBS before aliquoting the culture and trapping the bacteria in agarose pads containing 7H9 medium supplemented with ADC and Glycerol. The agarose pads were placed at 37 °C for 3 h to let the bacteria kickstart their metabolism. After 3 h, the pads were imaged under 63X objective of Leica SP8 (Leica Microsystems) under RFP and GFP channels. The total bacteria expressing RFP signal as well as GFP signal were quantified.

### Genomic and phage DNA isolation and estimation of phage DNA

DNA isolation was performed by Phenol-Chloroform-Isoamyl alcohol (PCI) extraction method. Briefly, 1 mL of culture aliquot was resuspended in 400 µL of TGE buffer (Tris-Glucose-EDTA Buffer) supplemented with 70 µL of 1 mg/mL Lysozyme and incubated overnight at 37 °C. Post incubation, 20 µg of proteinase K, 0.5% SDS, and 20 mM EDTA were added and incubated at 60 °C for 2 h. Vials were placed on ice and 500 µL PCI (25:24:1) was added and mixed rigorously prior to centrifugation at 14000 rpm, at 4 °C for 10 mins. Upper aqueous layer was collected and equal volume of Chloroform: Isoamyl alcohol (49:1) was added and centrifuged at 14000 rpm at 4 °C for 10 mins. Upper aqueous layer was collected, and 0.6 V isopropanol and 0.1 V 3 M sodium acetate was added and incubated on ice for 1 h. The tubes were then spun at 14000 rpm at 4 °C for 30 min and supernatant was discarded. The DNA pellet was washed with 500 µL of chilled 70% ethanol and resuspended in 50 µL of Milli-Q water. DNA concentration was determined using a nanodrop.

To estimate the phage DNA extracted, qPCR was performed. Phage specific primers were used to quantify the total amount of bacteriophage DNA. Circularization primers were used to determine the circularity of the phage DNA. SigA specific primers were used to determine the quantity of bacterial DNA for *M. smegmatis*. RpoD specific primers were used to determine the quantity of bacterial DNA for *P. aeruginosa*. Each PCR reaction (20□ μL) contained 10□ μL of 2×SYBR Green Master Mix Reagent (Applied Biosystems), 2 ng/μl of total genomic DNA and 200□ nM genome-specific primers. The reactions were performed in an RT-PCR machine (CFX96, Bio-Rad, US). The thermocycling conditions were 95 °C for 5□ min and 40 cycles at 95 °C for 30 s, 60 °C for 30□ s and 72 °C for 30□ s. Amplification specificity was assessed using melting curve analysis. Relative abundance of phage genome is calculated by normalizing Quantification Cycle (Cq) values of phage DNA to those of bacterial genome. Phage circularity was calculated as a percentage of Cq values of circularization specific-primer reactions to Cq values of phage genome specific primer reactions.

### Single cell growth assay

For *M. smegmatis* and *P. aeruginosa*, log phase bacterial cultures were resuspended in PBS for 48 h and were phage exposed for 3 h followed by AFS treatment to remove extracellular phages. Cultures were incubated at 37 °C, 180 RPM and at periodic intervals (from day 0 post infection to day 35 post infection for *M. smegmatis* and day 7 post infection for *P. aeruginosa*), aliquots were drawn, and single cell suspension was made by sonicating the aliquots three times-15 s ON and 15 s rest. Bacterial cultures were then passed through 26G needle at least 10 times before centrifuging at 300 g for 5 mins. Supernatant was collected and OD was measured before diluting the culture to low dilutions of 1000 CFU/mL, 100 CFU/mL and 10 CFU/mL. 10 µL of the three cultures was seeded into 90 µL of growth media in a 96-well plate well. The outer wells were filled with 200 µL autoclaved milliQ water. Plates were incubated at 37 °C for 7 days. At 7 days, OD measurements were taken along with visual count of wells showing growth. A cutoff of OD 0.5 was set to determine the number of wells showing growth. Dilutions showing 30-40% growth in non-phage exposed samples were considered and bacterial growth in phage exposed samples was normalized to non-phage exposed samples to obtain Relative Establishment Probability (REP)^48^.

For *M. tuberculosis,* log phase bacterial cultures were resuspended in PBS for 7 d to induce non-replication. Phages were exposed for 3 h followed by AFS treatment to remove extracellular phages. Cultures were then incubated at 37 °C, 180 RPM and at periodic intervals (from day 0 post infection to day 115 post infection), aliquots were drawn, and single cell suspension was made as described above. Supernatant was collected and OD was measured before diluting the culture to low dilutions of 1000 CFU/mL, 100 CFU/mL and 10 CFU/mL. 10 µL of the three cultures was seeded into 90 µL of growth media in a 96-well plate well. The outer wells were filled with 200 µL autoclaved milliQ water. Plates were incubated at 37 °C for 5 weeks with weekly replenishing of outer wells. After 5 weeks, the wells were visually counted for growth as the lack of tween and shaking resulted in *M. tuberculosis* growing as microaggregates. Dilutions showing 30-40% growth in non-phage exposed samples were considered and bacterial growth in phage exposed samples was normalized to non-phage exposed samples to obtain REP.

### Resazurin assay under non-replicating conditions

Metabolic activity of non-replicating *M. smegmatis* was quantified using resazurin assay. Briefly, 48 h PBS starved log phase *M. smegmatis* cultures were exposed to phages (TM4 or D29) at 10:1 MOI for 3 h. AFS treatment was carried out to remove extracellular phages and prevent further phage infection. The cultures were incubated at 37 °C and 180 RPM. At regular intervals, cultures were aliquoted and 2×10^7^ bacterial cells were seeded per well in a 96-well plate, in PBS. 30 µL of 0.01% resazurin, made in 20% tween-20, was added to each well and the plates were incubated at 37 °C for 24 h. Fluorescence was measured post incubation with an excitation at 530 nm and emission reading at 590 nm.

### Low-pH and antibiotic driven persistence models

For low-pH non-replication model, log phase secondary cultures of *M. smegmatis* were centrifuged at 5000rpm for 5 mins and resuspended in fresh filter-sterilized 7H9 supplemented with ADC and Glycerol, pH adjusted to 4.5 using 1N HCl. 0.1M MES hydrate buffer was added to maintain the pH at 4.5. Following resuspension, periodic CFU enumeration of clock plasmid containing *M. smegmatis* was carried out as described above to quantify the percentage of non-replicating cells. For phage efficacy experiments, TM4 was added at 10:1 MOI for 3h and extracellular phage was removed by AFS treatment and 3X centrifugation following which the cultures were resuspended in pH 4.5 7H9 growth media. Single cell growth assay was performed as described above.

For antibiotic-driven persistence model, log phase secondary cultures of *M. smegmatis* were treated with a combination of rifampicin, isoniazid and ethambutol at 2X MIC (30 µg/mL for rifampicin, 20 µg/mL for isoniazid and 1 µg/mL for ethambutol). Periodic sampling and CFU enumeration was carried out to quantify non-replication. For phage efficacy experiments, TM4 was added at 10:1 MOI for 3h and extracellular phage was removed by AFS treatment and 3X centrifugation following which the cultures were resuspended in antibiotic containing 7H9 growth media. Single cell growth assay was performed as described above.

For antibiotic-driven persistence model in *P. aeruginosa*, log phase secondary cultures were resuspended in complete M9 media and treated with a combination of meropenem and tobramycin at 10X MIC (7.5 µg/mL) and 5X MIC (3.75 µg/mL) respectively. Periodic sampling and CFU enumeration was carried out to quantify non-replication. For phage efficacy experiments, cultures treated with meropenem and tobramycin at 10X MIC and 5X MIC respectively for 24 h were exposed to DMS3vir-acrF1 at 10:1 MOI for 3h and extracellular phage was removed by 3X centrifugation following which the cultures were resuspended in antibiotic containing M9 complete media. Single cell growth assay was performed as described above.

### Single cell microscopy

Log phase bacterial cultures that have been PBS starved for 48 h were exposed to phage for 3 h prior to AFS treatment. Single cell suspension was obtained as described previously. Agarose pads were prepared by mixing (1:1) 1.5% agarose and autoclaved 2X 7H9 growth media supplemented with Glycerol and ADC and pouring onto glass coverslips with 45 min set time. Single cell suspension was then poured onto the flat surface of agarose pads and allowed to set for 30 mins post which, the pads were transferred to 35 mm imaging dishes. The pads were imaged with Leica SP8 confocal microscope (Leica Microsystems) with an imaging frequency of 1 h for timelapse microscopy for a total period of 24 h at 37 °C. The imaging chamber was humidified to minimize agarose shrinking over the period of 24 h. A custom python script was used to segment and quantify the cells, binning them into either dividing cells, dying cells or cells remaining in stasis. Briefly, the z-stack timelapse images were aligned through linear stack alignment with SIFT using ImageJ. Aligned images were segmented by applying a 3 x 3 Sobel edge detection filter, followed by thresholding using isodata thresholding algorithm. The resulting edges were skeletonized using Zhang and Suen algorithm. Cell areas were reconstructed from the skeletons. These cell area binary masks were used to label and track cells through frames by overlaying previous segmented frames. For quantification of cells into the three distinct phenotypes, the following methodology was used – For a given frame n; x, y position of cell i was recorded and a nearest neighbour search was performed (cell j). Nearest neighbour search was performed for n+5 (5 h). If a new nearest neighbour is detected, cell label was verified. If no cell label was found, a birth event was recorded and cell i was removed following the assignment of two new cell labels. For death events, a similar algorithm was followed but once a nearest neighbour (cell j) is detected, n-2 frames were searched for cell j. If cell j existed in all previous frames, a death event is recorded. Else, cell j is removed. Cells in stasis were calculated by subtracting the dividing and dying cells from the total number of cells in frame 1.

For *P. aeruginosa*, due to the motile nature of the bacteria, a higher percentage of agarose was used to make the pads. Briefly, 3% agarose was mixed with MilliQ water (1:1) and poured onto glass coverslips. Single cell suspension of nutrient starved *P. aeruginosa* was prepared as described previously and poured onto the flat surface of agarose pads and allowed to set for 30 mins before transferring to 35 mm imaging dishes. The pads were imaged with Leica SP8 confocal microscope (Leica Microsystems) with an imaging frequency of 30 min for timelapse microscopy for a total period of 24 h at 37 °C.

### SYBR Gold tagging of TM4 and FACS

In order to isolate only phage infected bacteria and study the single-cell growth, we tagged the TM4 bacteriophage with SYBR Gold dye. Briefly, 3X final concentration of SYBR Gold dye was added to the phage and incubated at 37 °C overnight. Post incubation, phage samples were centrifuging at 30,000 g for 3 h at 4 °C. Supernatant was discarded and the phage pellet was resuspended in phage buffer overnight at 4 °C. This process was continued for 3 times to remove free dye. SYBR Gold tagged phages were added to non-replicating *M. smegmatis* at 10:1 MOI for 3 h and extracellular phages were removed by washing three times with PBS and once with Ammonium Ferric Sulphate. Single cell suspension was obtained as previously described and samples were sorted (BD FACSAria Fusion, US) and high SYBR Gold positive cells were collected. Briefly, cells were first gated on forward scatter area (FSC-A) versus side scatter area (SSC-A) to exclude debris. Doublets were excluded using FSC-A versus FSC-H gating. SYBR Gold positive cells were identified as events with fluorescence intensity exceeding the highest signal from unstained control population. From the SYBR Gold-positive population, the top 44% to 51% highest GFP fluorescence intensity events were selected for sorting (**Supp Fig 17**). Once established, identical gates were applied to all samples. Sorted populations were collected into MCT tubes. These cells were then used for single cell growth assays as previously described.

### Pseudolysogeny in implant-associated bacteria

The implants used in this study were made up of Stainless Steel (SS316L), with dimensions of 5mm x 5mm x 1mm. Implants were incubated in Piranha solution for 24 h to remove organic debris from the surface. They were then washed with MilliQ water (3x) and autoclaved prior to usage. Cultures of *Pseudomonas aeruginosa* were grown to a late log phase and made into aliquots of 1 mL each. To induce bacterial attachment to the surface of the implant, the implants were placed in these aliquots for 2 h at 37 °C, without shaking. To induce nutrient-limited non-replication, the implants were then taken out and placed in 200 μL of M9 salts (Himedia, Maharashtra, India) for 48 h at 37 °C, without shaking. Then, implants were incubated with DMS3vir-acrF1 (10:1 MOI), Meropenem (1XMIC) (0.75 μg/ml), Tobramycin (1XMIC)(0.75 μg/ml), both DMS3vir-acrF1 (10:1 MOI) and Tobramycin (1XMIC) or both DMS3vir-acrF1 (10:1 MOI) and Meropenem (1XMIC) in a volume of 200 μL. To facilitate phage adsorption and infection, 2 μL of sterile 0.1M CaCl2 was supplemented to each treatment solution. This treatment lasted for 24 h, at 37 °C, without shaking. The implants were then taken out of the treatment solutions and placed in 200 μL of M9 (Himedia, Maharashtra, India) salts for 24 h, at 37 °C, without shaking. The implants were then sonicated for 90 s in a bath sonicator to suspend all implant associated bacteria. The solutions were then serially diluted in a 96-well plate and spotted on LB-agar plates to calculate CFUs.

### Protospacer acquisition check

To check protospacer acquisition in *P. aeruginosa,* log phase secondary cultures were washed and resuspended in PBS. 48 hours post nutrient starvation, cultures were treated with DMS3vir at 10:1 MOI for 3 h. Extracellular phage was removed by AFS treatment and 3X washes. Cultures were incubated at 37 °C for 7 days following which DNA was extracted as described previously. To assess the protospacer acquisition, qPCR was performed to amplify the flanking regions of protospacer acquisition region. Each PCR reaction (20□ μL) contained 10□ μL of 2×SYBR Green Master Mix Reagent (Applied Biosystems), 2 ng/μl of total genomic DNA and 200□ nM genome-specific primers. The reactions were performed in an RT-PCR machine (CFX96, Bio-Rad, US). The thermocycling conditions were 95 °C for 5□ min and 40 cycles at 95 °C for 30 s, 64 °C for 30□ s and 72 °C for 30□ s. Post PCR, samples were run on 3% agarose gel to visualize the length of amplicons in phage exposed vs non-exposed samples.

### *In vivo* implant associated non-replicating *Pseudomonas* model

Phage preparation: The bacterial lysate containing phages was centrifuged at 30,000g for 3 hr to pellet the phages. The phage pellet was resuspended in fresh phage storage buffer. The sample was then run through a QA-CIM monolithic column (BIA separations, Slovenia) using Fast Protein Liquid Chromatography (FPLC)(Biorad, USA)^85^. Briefly, sample was washed with 20% NaCl followed by 70% NaCl sample elution. Following FPLC, the purified phage was filtered and titred prior to animal studies.

For the pre-treatment studies, cultures of *Pseudomonas aeruginosa* were grown to a late log phase and PBS starved for 48 h following which they were exposed to DMS3vir-acrF1 for 3 h. Post 3 h, extracellular phage was washed away by treating with Ammonium Ferric Sulphate and centrifugation. The bacteria were then allowed to attach to SS316L implants for 24 h following which the implants were surgically placed into Balb/c mice subcutaneously. Post surgery, animals were administered Meloxicam (5mg/kg/day) to alleviate pain and inflammation. Animal weights were monitored until the end of the experiment. 48 h post implantation, animals were sacrificed and the implants were recovered and suspended in PBS. Sonication for 90 s was carried out to detach the bacteria from the implant surface and the bacteria was enumerated through CFU assay.

For treatment studies, cultures of *P. aeruginosa* were grown to a late log phase and PBS starved for 48 h following which they were allowed to attach to implants for 2 h in PBS. Post attachment, implants were surgically placed into C57BL/6 mice subcutaneously. A different strain of mice was used to rule out strain specific outcomes and increase rigour of the experiment. Post surgery, animals were administered Meloxicam (5 mg/kg/day) subcutaneously to alleviate pain and inflammation along with either saline, phage (10:1 MOI), meropenem (300 mg/kg/day) or a combination of phage (10:1 MOI) and meropenem (300 mg/kg/day) subcutaneously at the site of surgery. The experiment was terminated 24 hours post-surgery due to mortality in saline treated group and the implants were extracted to enumerate the CFUs.

For antibiotic induced persistence studies, cultures of *P. aeruginosa* were grown to a late log phase and PBS starved for 48 h following which they were allowed to attach to implants for 2 h. 24 h prior to surgery, animals were administered Tobramycin (3 mg/kg/day, 6 mg/kg/day and 12 mg/kg/day for optimization, 12 mg/kg/day for checking CRISPR activity) intraperitoneally. Following bacterial attachment, implants were surgically placed into C57BL/6 mice subcutaneously. Animals were administered Meloxicam (5 mg/kg/day) subcutaneously immediately and another dose at 24 h post-surgery to alleviate pain and inflammation along with either saline, DMS3vir (10:1 MOI), or DMS3vir-acrF1 (10:1 MOI) subcutaneously near the surgery site. Mice were administered Tobramycin (12 mg/kg/day) intraperitoneally on the day of the surgery and the following day. The experiment was terminated 48 hours post-surgery and the implants were extracted to enumerate the CFUs.

### Statistical Analysis

All experiments were performed on independent biological replicates. Mean and standard deviation are reported in the figures. Statistical significance was determined for control and experimental groups using multiple unpaired two-sided t-test with alpha = 0.05. For analysis of multiple groups, 1-way ANOVA with Tukey test or 2-way ANOVA with Sidak’s multiple comparisons test with alpha = 0.05 were used. GraphPad (Prism) was used for all statistical analysis.

## Data Availability

The data generated in this study has been deposited in the public database:

RNA sequence raw files along with reference genomes and GTF files are deposited in the public repository:

## Code Availability

Code used to analyze and quantify single cell tracking microscopy data is deposited in Github: https://github.com/yeswanthck/Single-Cell-Tracking-Analysis

## Ethical Approvals

This study was approved by the Institutional BioSafety Committee (IBSC) under protocol numbers IBSC/IISc/RA/14/2024 and IBSC/IISc/RA/03/2024 for *P. aeruginosa* and IBSC/IISc/RA/19/2023-24 for *M. tuberculosis*. The animal experiments were performed following the Institute Animal Ethics Committee (IAEC) protocols CAF/Ethics/070/2024 and CAF/Ethics/123/2025.

## Notes

### Competing Interest Statement

The authors have declared no competing interest.

