## Supplemental Information for "Bacteriophages utilize pseudolysogeny to target non-replicating bacteria and CRISPR-resistant phages eliminate recalcitrant implant infections"

#### Supplementary Information

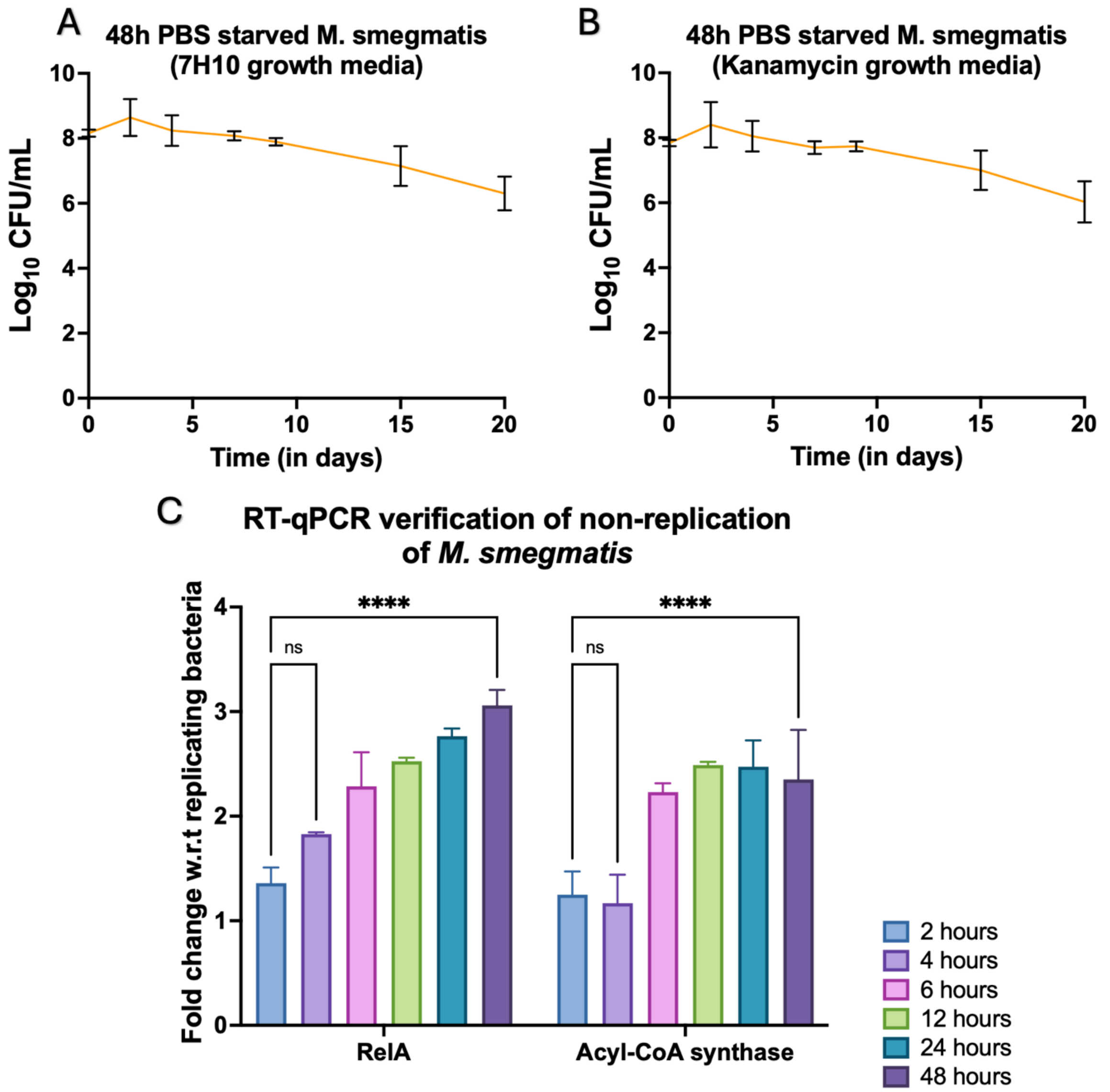


***Supplementary Figure 1. Verification of M. smegmatis non-replication****. Enumeration of 48h 1X PBS starved M. smegmatis containing clock plasmid on A) 7H10 growth medium B) Kanamycin containing 7H10 growth medium to enumerate plasmid containing M. smegmatis C) Gene expression levels of RelA (left) and Acyl-CoA (right) synthase in nutrient starved M. smegmatis. (n = 3) Two way ANOVA with Šídák's multiple comparisons test was used to determine the statistical significance. (*p ≤0.05, **p ≤0.01, ***p ≤0.001, ****p ≤0.0001).*

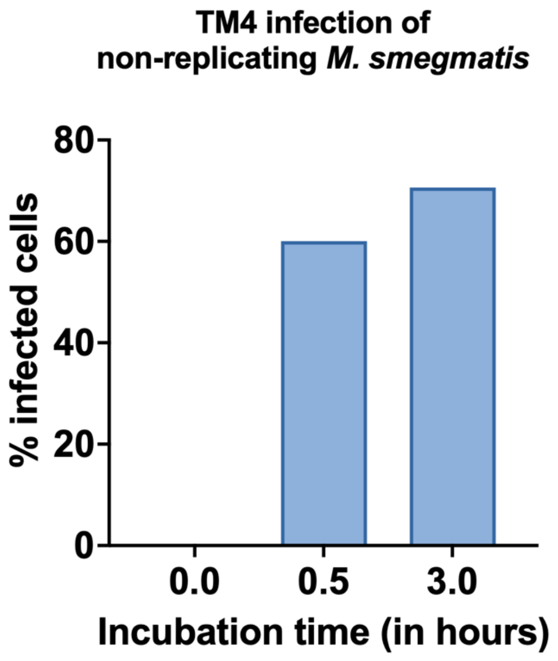


***Supplementary Figure 2. Verification of TM4 phage infection against non-replicating M. smegmatis***. *Phage infection quantification in non-replicating M. smegmatis through SYTO9 tagged TM4 bacteriophage (n=1).*

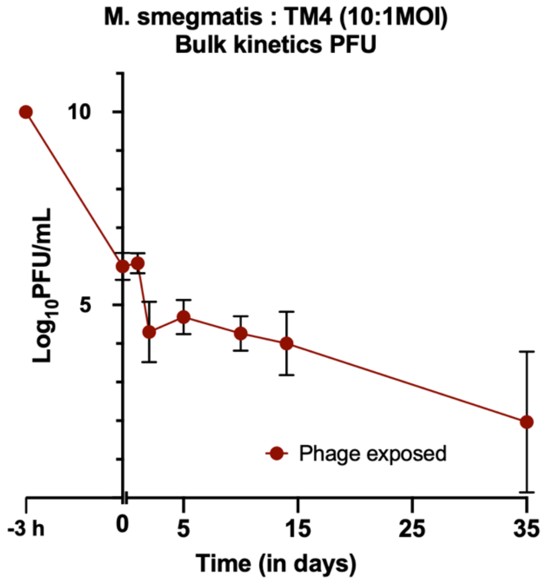


***Supplementary Figure 3. PFU kinetics of TM4 exposed non-replicating M. smegmatis***. *Plaque forming units (PFUs) of extracellular bacteriophages in the supernatant (n=3).*

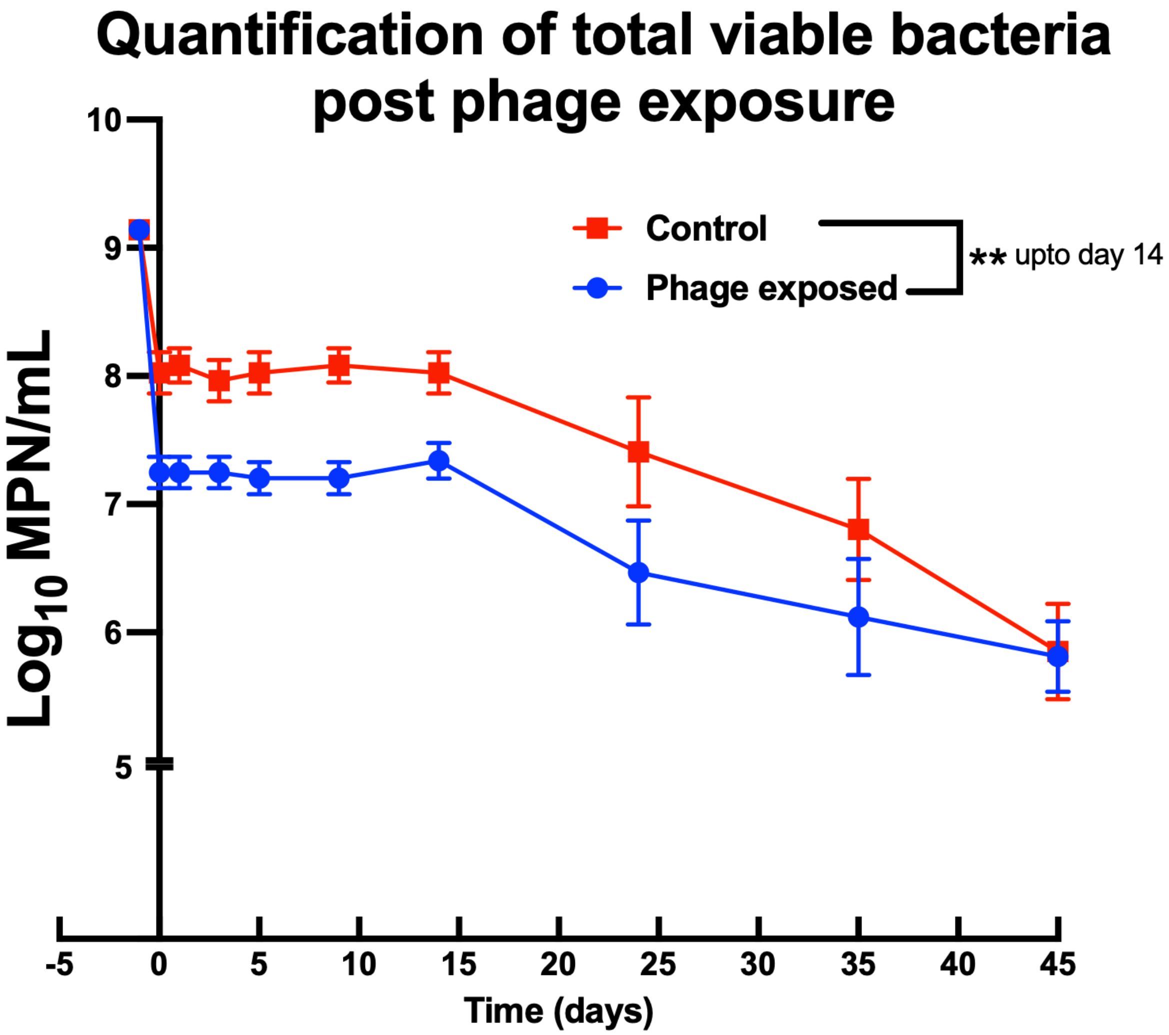


***Supplementary Figure 4. Most probable number enumeration.*** *MPN assay of non-replicating M. smegmatis with or without TM4 phage exposure at 10:1 MOI. (n=3) Two way ANOVA with Holm-Šídák's multiple comparisons test was used to determine the statistical significance. (*p ≤0.05, **p ≤0.01, ***p ≤0.001, ****p ≤0.0001).*

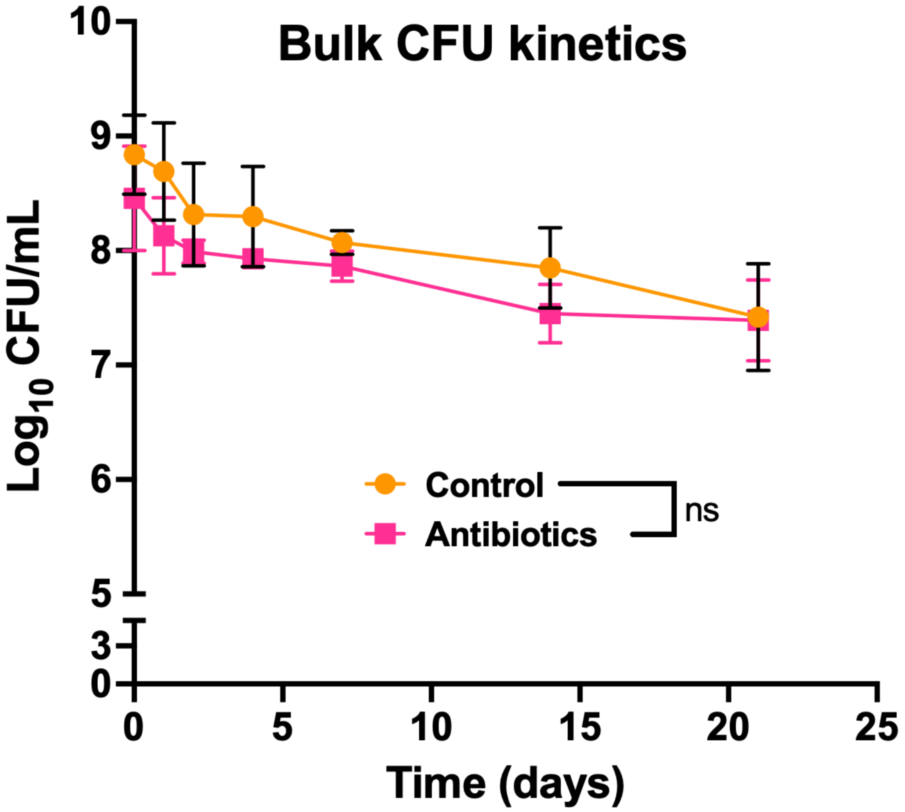


***Supplementary Figure 5. Bulk kinetics of Antibiotic treated non-replicating M. smegmatis***. *48 h PBS starved non-replicating M. smegmatis was treated with a combination of Rifampicin (30 µg/mL), Isoniazid (20 µg/mL), and Ethambutol (1 µg/mL) (n=3). Two way ANOVA with Holm-Šídák's multiple comparisons test was used to determine the statistical significance*

### Single cell microscopy quantification

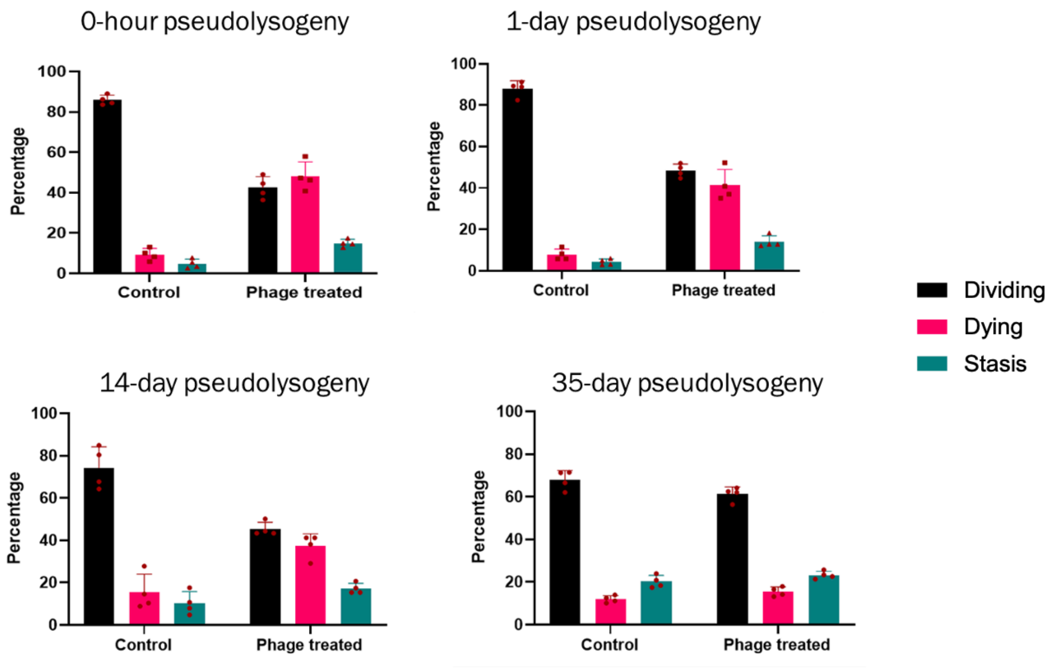


| Pseudolysogeny duration (in days) | Significance between control and phage exposed |
|---|---|
| 0 | Dividing - ****, Dying – ****, Stasis – ns, |
| 1 | Dividing - ****, Dying – ****, Stasis – *, |
| 14 | Dividing - ****, Dying – ***, Stasis – ns |
| 35 | Dividing - *, Dying – ns, Stasis – ns |

***Supplementary Figure 1***. *Quantification of single cell timelapse microscopy of non-replicating M. smegmatis exposed to TM4 bacteriophage for 3 hours prior to growth media supplementation after several days. (n= 4) Two way ANOVA with Holm-Šídák's multiple comparisons test was used to determine the statistical significance. (*p ≤0.05, **p ≤0.01, ***p ≤0.001, ****p ≤0.0001).*

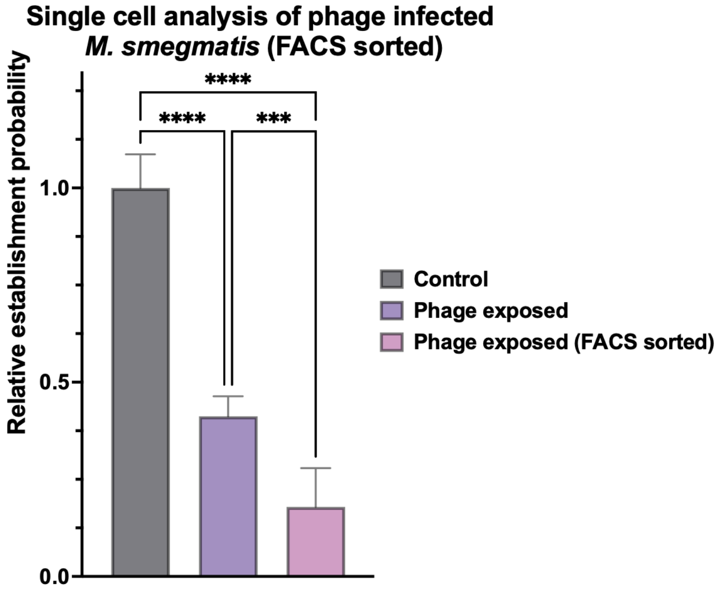


***Supplementary figure 7. Single cell growth analysis of TM4 exposed M. smegmatis*** *A) SYBR Gold-tagged TM4 phage efficacy towards phage infected bacteria only quantified through single-cell growth assay of FACS sorted M. smegmatis (Data pooled from two independent experiments). (n = 3) One-way ANOVA was used to determine statistical significance. (*p ≤0.05, **p ≤0.01, ***p ≤0.001, ****p ≤0.0001).*

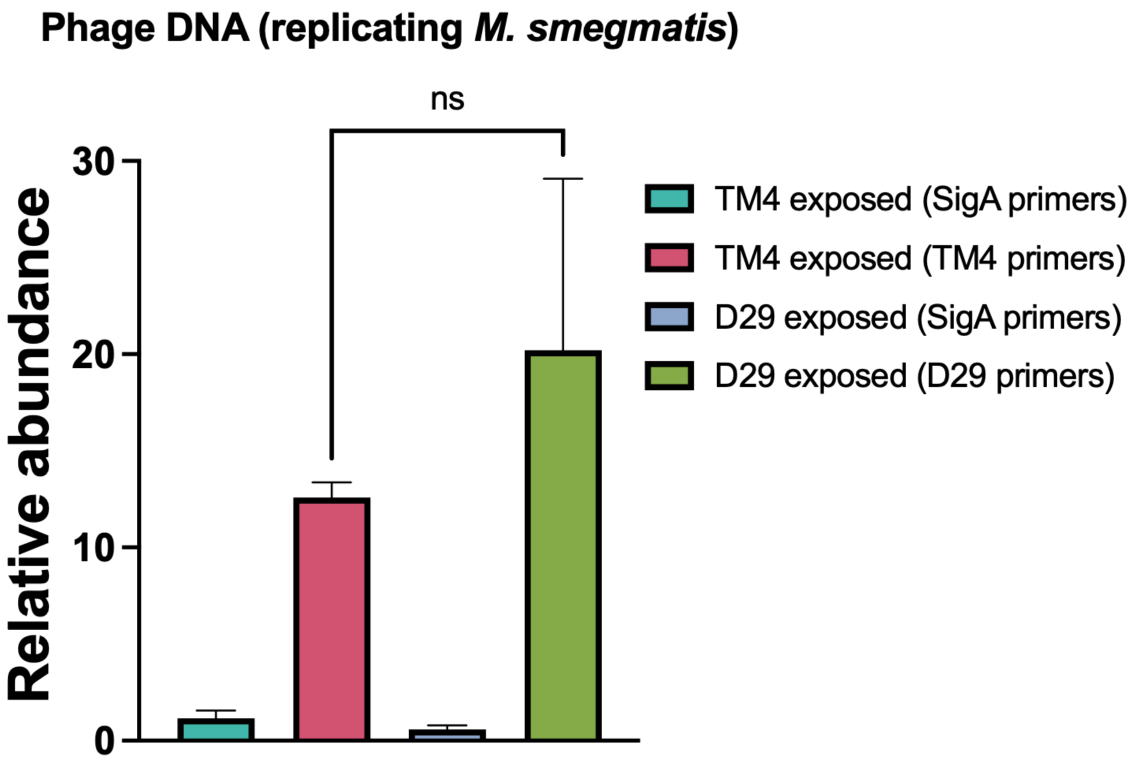


***Supplementary Figure 8.*** *DNA quantification in replicating M. smegmatis. DNA was extracted 20 mins post phage infection in respective groups. Unpaired t-test was used to test statistical significance (n = 3) (*p ≤0.05, **p ≤0.01, ***p ≤0.001, ****p ≤0.0001).*

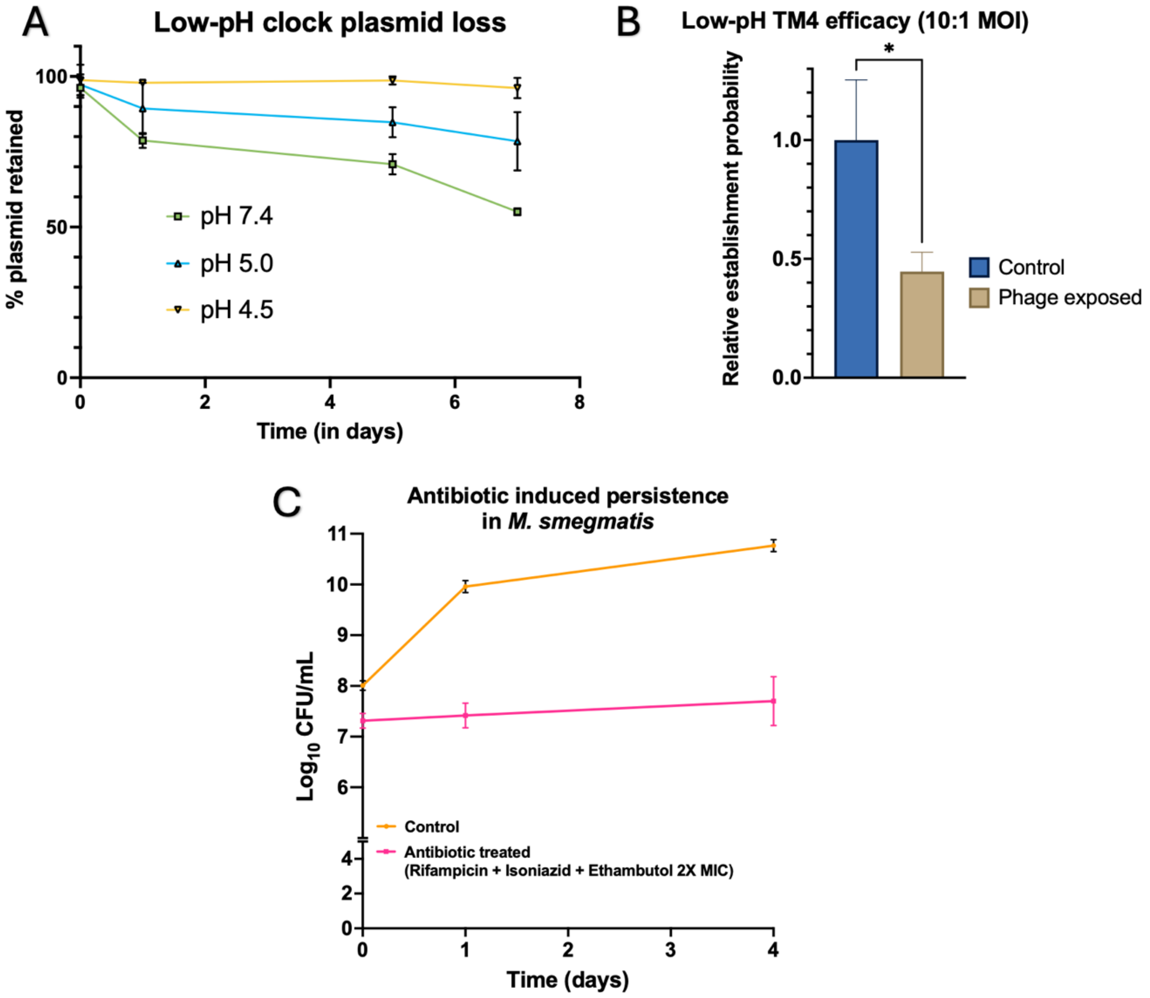


***Supplementary Figure 9. Low-pH and antibiotic-induced non-replication model of M. smegmatis***
***A)*** *Percentage of surviving bacteria with the clock plasmid retained (n=3)* ***B)*** *Phage efficacy quantification on low-pH (pH 4.5) non-replicating M. smegmatis (n=3)* ***C)*** *Antibiotic induced persistence of M. smegmatis when treated with a combination of Rifampicin, Isoniazid, and Ethambutol at 2X MIC (n=3). For* ***B****, one way ANOVA was used to determine the statistical significance. (n= 3) (*p ≤0.05, **p ≤0.01, ***p ≤0.001, ****p ≤0.0001).*

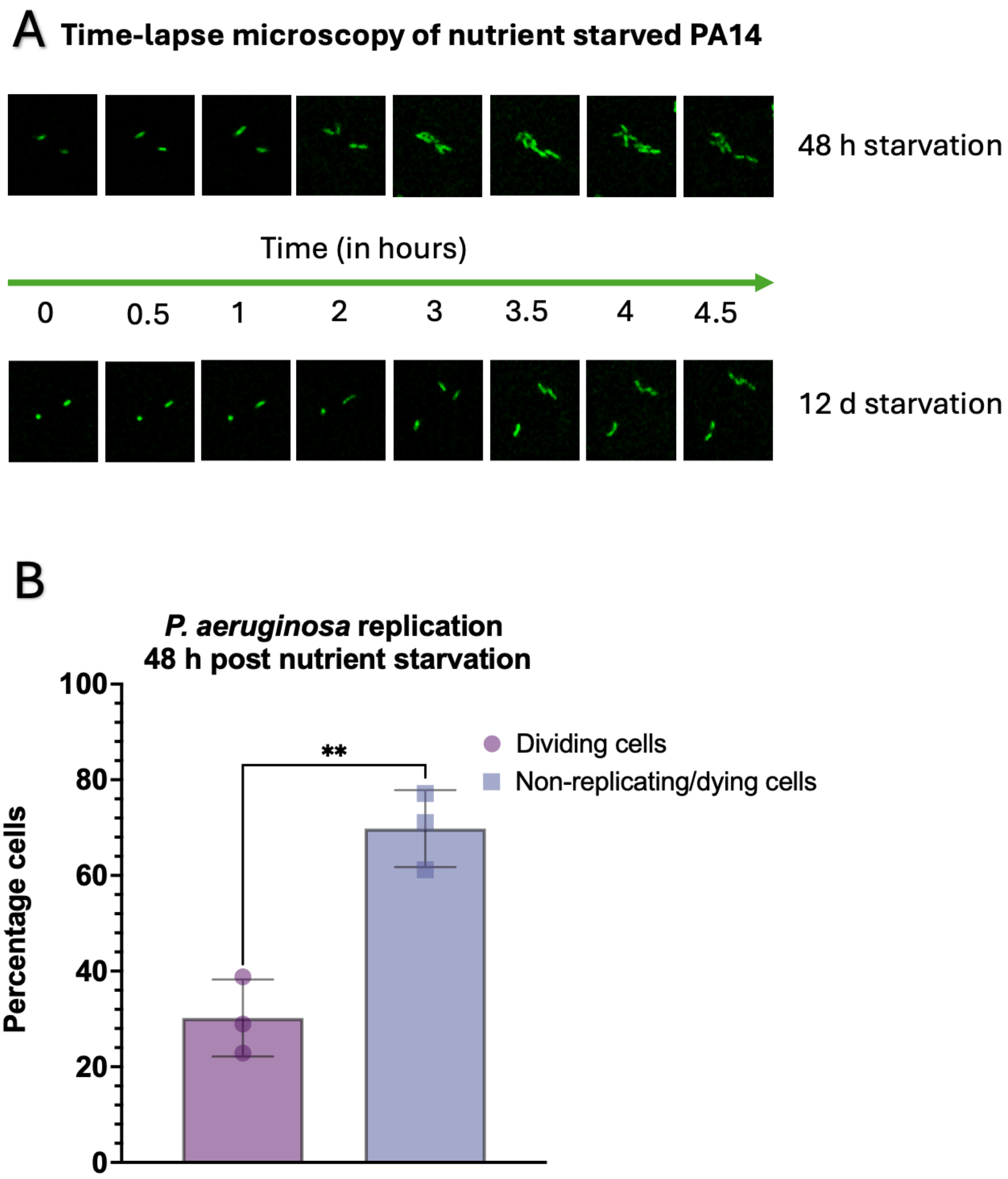


***Supplementary Figure 10. Replication of nutrient stressed P. aeruginosa. A)*** *Timelapse microscopy of 48 h PBS starved (top) and 12 day PBS starved (bottom) P. aeruginosa reveals the presence of replicating bacterial cells.* ***B)*** *Quantification of single-cell microscopy of 48 h PBS starved P. aeruginosa for percentage of replicating cells and non-replicating cells. Unpaired t-test was used to test statistical significance (n = 3) (*p ≤0.05, **p ≤0.01, ***p ≤0.001, ****p ≤0.0001).*

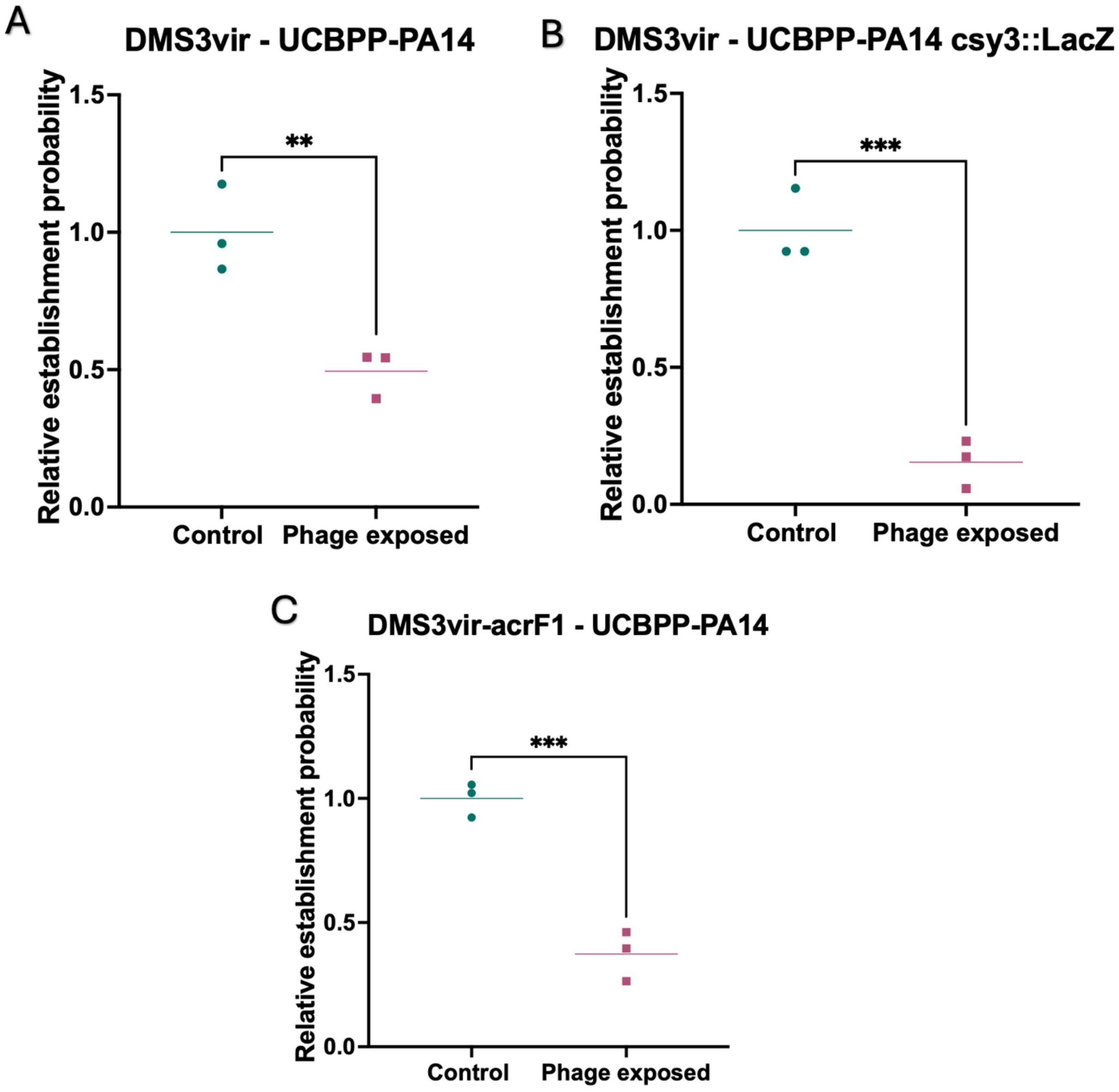


***Supplementary Figure 11. Phage efficacy against nutrient stressed P. aeruginosa A)*** *CRISPR susceptible DMS3vir phage infectivity and efficacy towards nutrient-stressed CRISPR-active P. aeruginosa quantified through single-cell growth assay (n=3)* ***B)*** *CRISPR susceptible DMS3vir phage infectivity and efficacy towards nutrient-stressed CRISPR-inactive P. aeruginosa quantified through single-cell growth assay (n=3)* ***C)*** *CRISPR resistant DMS3vir-AcrF1 phage infectivity and efficacy towards nutrient-stressed CRISPR-active P. aeruginosa quantified through single-cell growth assay (n=3). Unpaired t-test was used to determine the statistical significance (*p ≤0.05, **p ≤0.01, ***p ≤0.001, ****p ≤0.0001).*

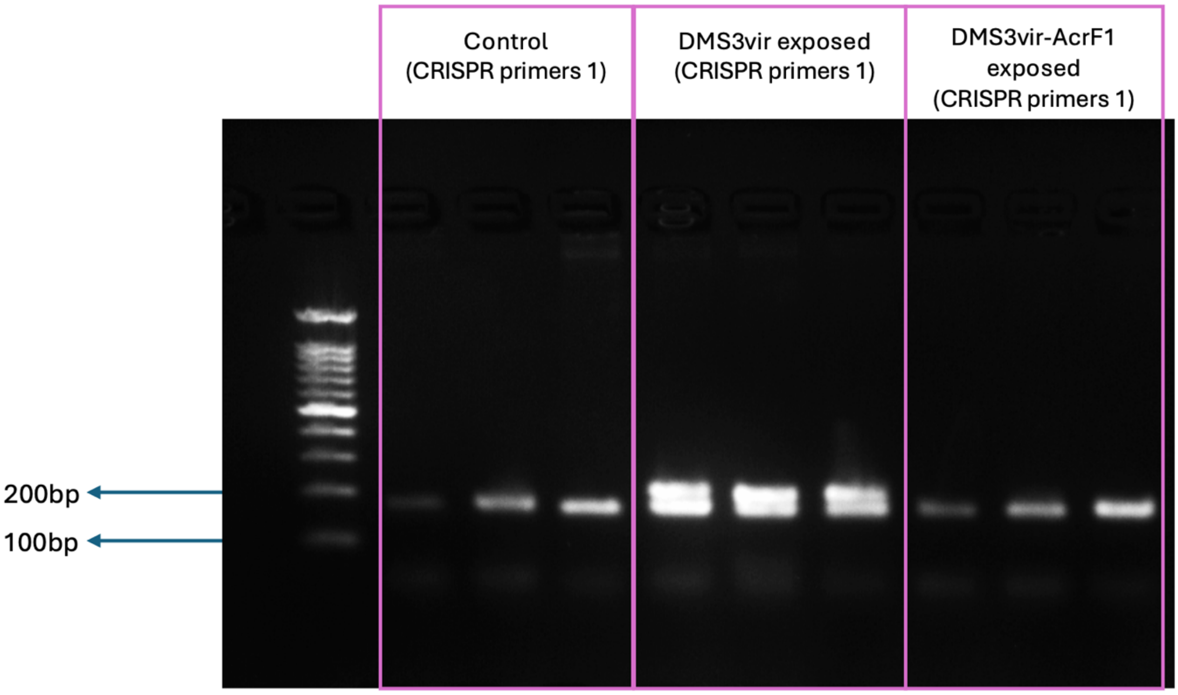


***Supplementary Figure 12. Protospacer acquisition***. *Gel electrophoresis post RT-PCR for UCBPP-PA14 DNA showing acquisition (increase in the amplicon size) 1 day post exposure to DMS3vir or DMS3vir-AcrF1 at 10:1 MOI (n=3).*

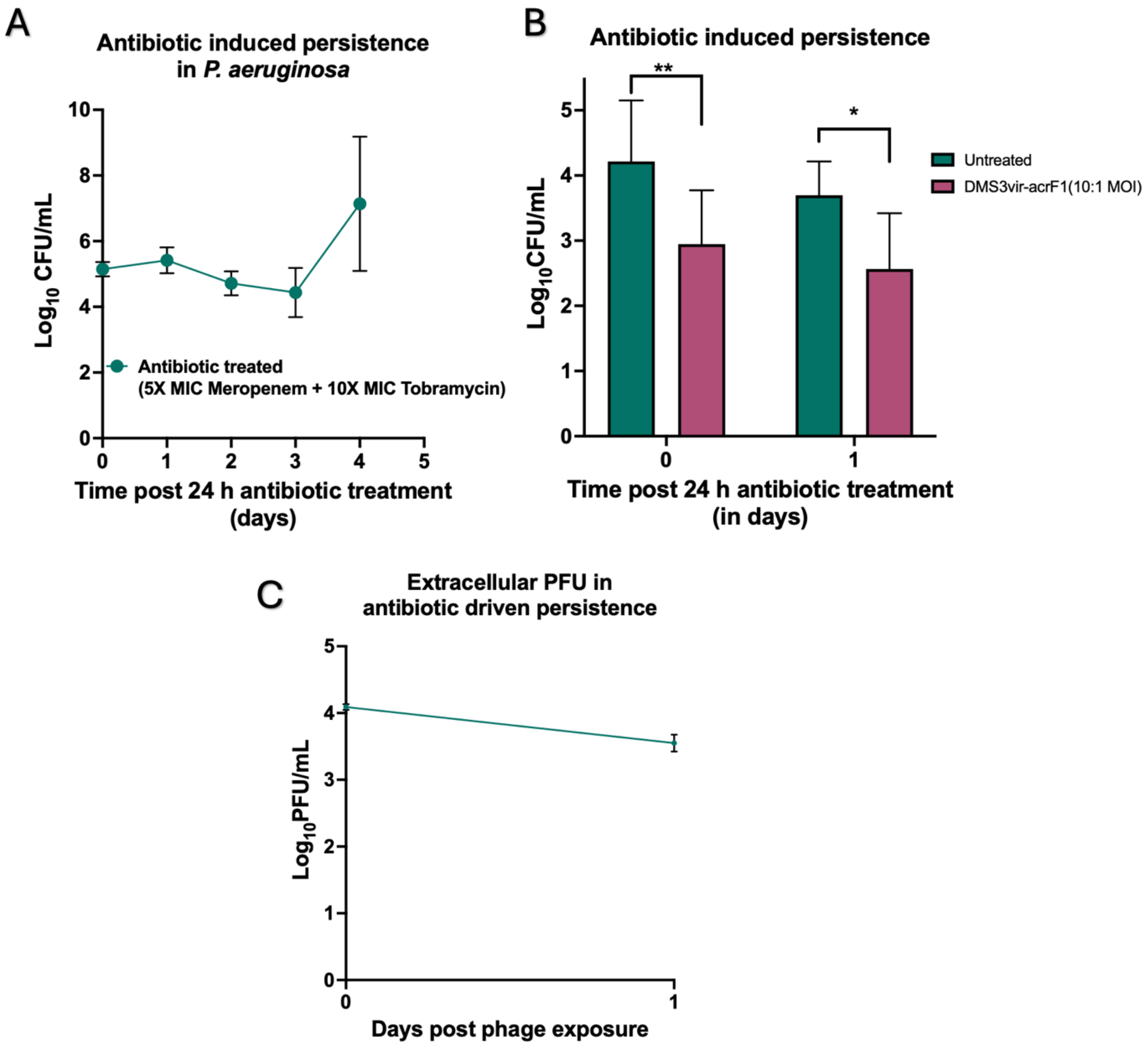


***Supplementary Figure 13. Antibiotic induced persistence in Pseudomonas aeruginosa. A)*** *Antibiotic induced persistence in P. aeruginosa over multiple days when treated with a combination of 5X MIC meropenem and 10X MIC tobramycin (n=3).* ***B)*** *Phage efficacy against antibiotic induced persistence in P. aeruginosa (n=3).* ***C)*** *Extracellular PFUs in antibiotic induced persistence model showing lack of phage replication until nutrient supply and antibiotic stress removal (n=3). For B, Two way ANOVA with Holm-Šídák's multiple comparisons test was used to determine the statistical significance. (n= 3) (*p ≤0.05, **p ≤0.01, ***p ≤0.001, ****p ≤0.0001).*

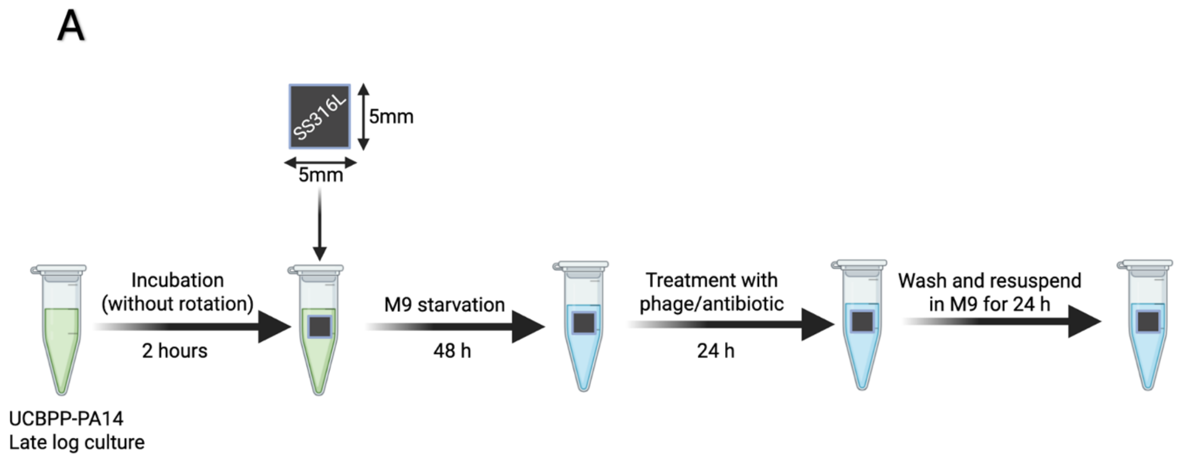


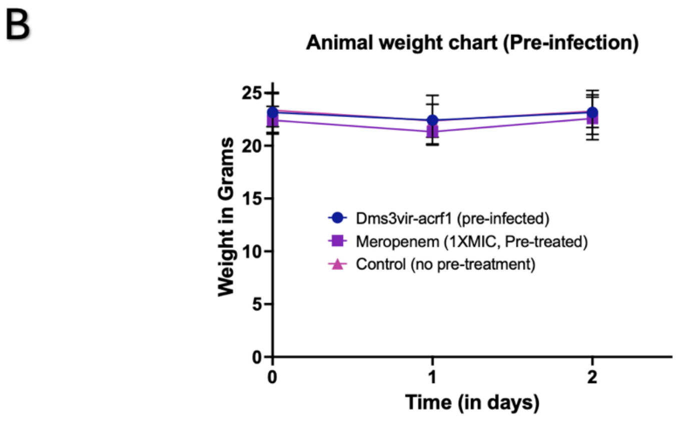


***Supplementary Figure 14. Implant associated nutrient stressed P. aeruginosa model*** **A)** *Schematic representation for obtaining implant-associated non-replicating P. aeruginosa.* **B)** *Weight chart of Balb/c mice post implantation of P. aeruginosa harbouring stainless steel SS316L implants (n=10, data pooled from two individual experiments).*

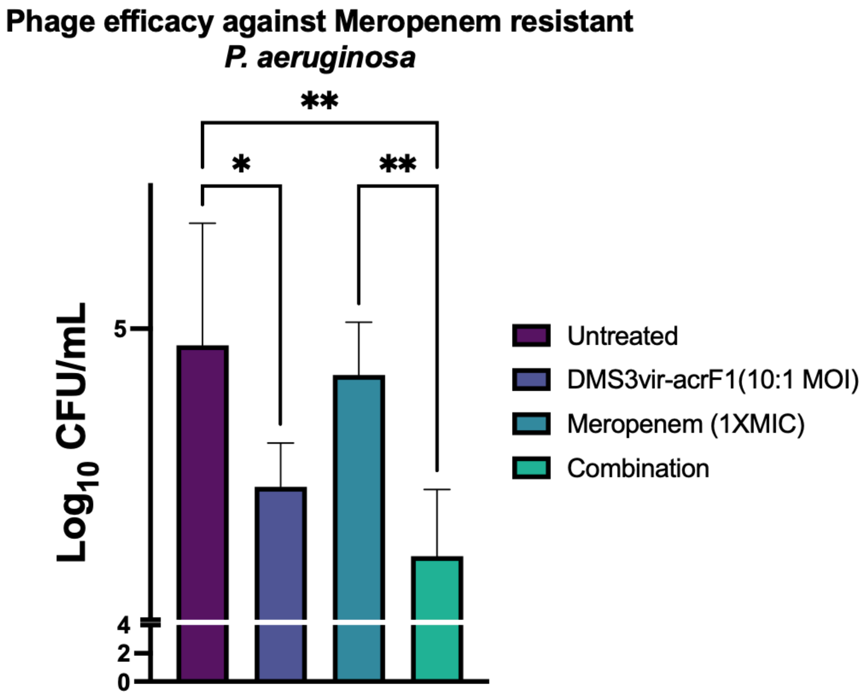


***Supplementary Figure 15. Phage efficacy against MDR-strain of P. aeruginosa.*** *DMS3vir-acrF1 efficacy against implant associated nutrient-stressed multidrug resistant clinical isolate of P. aeruginosa BEI 12368. One-way ANOVA was used to determine the statistical significance (n=3). (*p ≤0.05, **p ≤0.01, ***p ≤0.001, ****p ≤0.0001).*

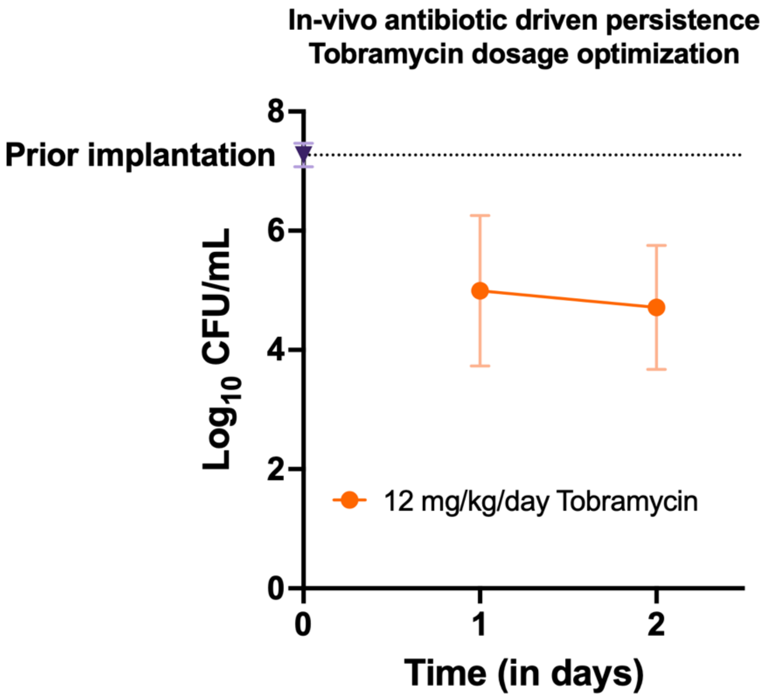


***Supplementary Figure 16. In vivo antibiotic induced persistence of implant associated P. aeruginosa.*** *Optimization of Tobramycin concentration to ensure stable CFU post initial drop. Mice were administered 12 mg/kg/day of tobramycin intraperitonially on Days -1, 0 and 1 (n=5). Dotted line represents in vitro CFU on the implant prior to implantation.*

A

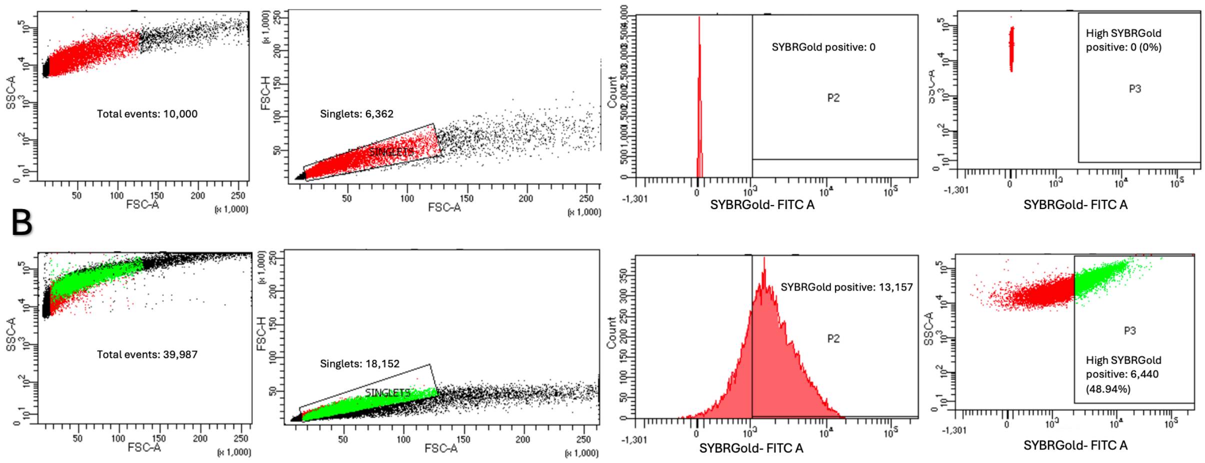


B

***Supplementary Figure 17. Gating strategy for flow-sorting of fluorescently tagged phage infected M. smegmatis.*** *A) Unstained control used to draw gates for SYBR Gold-FITC A channel excluding 100% of unstained cells B) Fluorescently tagged phage infected bacterial sorting. Cells were first gated on FSC-A versus SSC-A to exclude debris. Singlets were selected using FSC-A versus FSC-H gating. SYBR Gold positive cells were identified as events with fluorescence intensity exceeding the highest signal from unstained control population. From the SYBR Gold-positive population, the top 44% to 51% highest GFP fluorescence intensity events were collected and single-cell growth assay was performed.*

***Table S1.* List of primers used in this study.**

| Primer name | Forward sequence (5'-3') | Reverse sequence (5'-3') | Purpose |
|---|---|---|---|
| RelA | TTCCCGCCACCAAAGGCAAT | AATATTGCGCCCGGCCATGA | Quantification of RelA cDNA |
| AcSyn | GCTGACCGCGTCGAAGATCA | CGCGTAGCAGTCCCGGATTT | Quantification of Acyl CoA synthase cDNA |
| SigA | CGTTCCTCGACCTCATCCAG | TGATCACCTCGACCATGTGC | Housekeeping gene *M. smegmatis* |
| D29 | GTTTAGCGCCGCTGGCATAAG | CACACACCAATTTGCAATTCGG | Quantification of D29 phage DNA |
| TM4 | CGTTCGCTGTCGAGTTCTGTG | CGCATCGAGCACAGGTAGGTAC | Quantification of TM4 phage DNA |
| D29_Cos | TTTGCTGTGCATGCCATCGT | GATCGGCTTGGAAACCCGTA | Quantification of D29 phage DNA circularization |
| TM4_Cos | CGTTTGTGCAGGTCAGCGA | CGGTGTTTGCTGGTGGAAAAA | Quantification of TM4 phage DNA circularization |
| CRISPR Primers 1[86] | GAGGGTTTCTGGCGGGAA | GTCCAGAAGTCACCACCCG | *P. aeruginosa* Protospacer acquisition check |
